# Yeast PI(3,4,5)P_3_ controls vacuole fusion through Vam7 and membrane microdomain assembly

**DOI:** 10.1101/2025.08.01.668199

**Authors:** Chi Zhang, Jorge D. Calderin, Robert P. Sparks, Aliasgar Topiwalla, Vito Failla, Daniel Grudzien, Soroush Asadi, Lylia Gomez, Yupeng Li, Ved Shah, Krish T Patel, Jahnavi M. Karat, Charlie T. Knapp, Razeen Ahmed, Emad Tajkhorshid, Rutilio A. Fratti

**Affiliations:** Department of Biochemistry, University of Illinois Urbana-Champaign, Urbana, IL 61801, USA; Center for Biophysics and Quantitative Biology., University of Illinois Urbana-Champaign, Urbana, IL 61801, USA; Department of Chemistry & Chemical Biology, Northeastern University, Boston MA 02115, USA; Theoretical and Computational Biophysics Group, NIH Center for Macromolecular Modeling and Visualization, University of Illinois Urbana-Champaign, Urbana, IL 61801, USA; Beckman Institute for Advanced Science and Technology, University of Illinois Urbana-Champaign, Urbana, IL 61801, USA

**Keywords:** Grp1-PH, Ypt7, SNARE, PIP3, HOPS, Vps33, PTEN, Vps34

## Abstract

Membrane trafficking is regulated by the spatiotemporal distribution of phosphoinositides. The endolysosomal pathway is controlled by PI3P, PI(4,5)P_2_ and PI(3,5)P_2_, whereas a role for PI(3,4,5)P_3_ is less clear. We report that yeast produce PI(3,4,5)P_3_ through Vps34 activity. In vitro assays showed that dioctanoyl (C8) PI(3,4,5)P_3_, the PI(3,4,5)P_3_-binding domain Grp1-PH and the phosphatase PTEN blocked vacuole fusion. Fluorescence microscopy showed that PI(3,4,5)P_3_ was present at the plasma membrane and vacuoles, and that its detection was blocked by PTEN, C8-PI(3,4,5)P_3_, the Vps34 inhibitor SAR405 and a *VPS34* temperature sensitive mutation. In addition, minimizing PI(4,5)P_2_ as a substrate for Vps34 with a *MSS4* temperature sensitive mutation reduced PI(3,4,5)P_3_ levels. Importantly, PI(3,4,5)P_3_ was required for the vertex enrichment of Ypt7 and the HOPS subunit Vps33. Finally, we showed that the soluble SNARE Vam7 was retained in a PI(3,4,5)P_3_-dependent manner and that its displacement from membranes blocked trans-SNARE pairing. These results demonstrate that vacuolar PI(3,4,5)P_3_ coordinates vertex assembly and SNARE function.

**Summary:** *Saccharomyces cerevisiae* produces PI(3,4,5)P_3_ via Vps34 activity on vacuoles to regulate homotypic fusion. PI(3,4,5)P_3_ functions through controlling vertex microdomain assembly and retaining the soluble SNARE Vam7 on membranes to drive fusion.

## Introduction

Membrane trafficking and fusion are driven by a group of conserved regulatory proteins (e.g., SNAREs) and lipids with organelle specificity. Regulatory lipids are relatively low in abundance yet carry out critical aspects of membrane trafficking (Krauss and Haucke, 2007; Lemmon, 2008; Corvera et al., 1999; Balla, 2013). These lipids include phosphoinositides (PI), phosphatidic acid (PA), diacylglycerol (DAG), sterols, lysolipids, and sphingolipids (Balla, 2013; Lingwood and Simons, 2010; Tu-Sekine et al., 2015; Xie et al., 2015; Starr and Fratti, 2019; Zhang et al., 2026; Reese and Mayer, 2005). PIs are glycerophospholipids with an inositol head group that can be phosphorylated at the D-3, D-4 and D-5 positions to generate seven distinct lipids whose primary function is the binding of proteins with domains that recognize specific variants. Broadly speaking, different organelles are marked by a dominant PI. Endosomes are marked by PI3P and PI(3,5)P_2_, whereas the Golgi contains PI4P, and the plasma membrane is populated by PI(4,5)P_2_ and PI(3,4,5)P_3_. While concentrated at characteristic organelles, biologically significant amounts can traffic to different membranes where they continue to signal and control membrane function.

The yeast vacuole/lysosome collects regulatory lipids from several pathways and continues to modify PIs to control homotypic vacuole fusion, vacuole fission and autophagic flux (Starr and Fratti, 2019; Bonangelino et al., 2002; Posor et al., 2022). Homotypic vacuole fusion can be divided into six stages, each of which is driven by a spatiotemporal-specific mixture of regulatory lipids. In a stage we now call *pre-priming*, Sec18 is sequestered from inactive cis-SNARE complexes by PA (Sasser et al., 2012; Starr et al., 2016, 2019). Sec18 can transfer to cis-SNARE complexes upon the conversion of PA to DAG by the phosphatase Pah1. Once Sec18 can engage SNAREs via its adaptor protein Sec17, *priming* occurs through ATP hydrolysis, which is dependent on ergosterol and PI(4,5)P_2_ through an undefined mechanism (Kato and Wickner, 2001; Mayer et al., 2000). Vacuole *tethering* occurs through the interaction of the Rab Ypt7 and its effector tethering complex HOPS (homotypic fusion and protein sorting) between partner membranes (Price et al., 2000b; a; Seals et al., 2000; Zhang et al., 2024). Ypt7 recruitment and activation is carried out by the guanine exchange factor (GEF) Mon1-Ccz1 bound to PI3P (Lawrence et al., 2014; Cabrera et al., 2014), while HOPS itself can simultaneously bind several PIs (Stroupe et al., 2006). PI3P is also essential for binding the PX domain of the soluble Qc-SNARE Vam7 and the formation of SNARE complexes during the *docking* stage (Boeddinghaus et al., 2002; Fratti and Wickner, 2007). Between docking and full content mixing, vacuoles can undergo hemifusion where only the outer leaflets of vesicles mix while leaving luminal contents separated by the inner leaflets (Reese and Mayer, 2005; Reese et al., 2005). We and others have found that this transition requires DAG (Jun et al., 2004) and is sensitive to exogenously added PI(3,5)P_2_ and lysophosphatidylcholine (Miner et al., 2019; Reese and Mayer, 2005).

During the tethering stage, vacuoles in contact are drawn together to form a flattened disc of membranes called the boundary domain. At the edge of the boundary disc, where membranes come into contact, are vertex points. Fusion machinery proteins and lipids that are initially dispersed throughout the membrane become enriched at vertices to build microdomains that form a discontinuous series of lipid-ordered patches around the boundary called the vertex ring. These domains are enriched in SNAREs, Ypt7, HOPS, actin as well as PI3P, PI(4,5)P_2_, DAG and ergosterol (Fratti et al., 2004; Wang et al., 2002, 2003; Eitzen et al., 2002; Karunakaran et al., 2012). Fusion occurs at vertex domains, leading to the internalization of the boundary as intralumenal vesicles that get degraded as part of a controlled mechanism to downregulate select membrane transporters (Mattie et al., 2017; McNally et al., 2017). Disruption of vertex domain assembly negatively impacts subsequent events including trans-SNARE complex formation, hemifusion and full content mixing.

While multiple PIs are essential in the formation of functional vertex microdomains, roles for the remaining PIs – PI5P, PI(3,4)P_2_ and PI(3,4,5)P_3_ – have not been assigned in vacuole homeostasis. PI(3,4,5)P_3_ is one of the most studied PIs that is primarily present at the plasma membrane where it directs many signal transduction pathways (Riehle et al., 2013). PI(3,4,5)P_3_ is made by class-I PI 3-kinase (PI3K) p110 isoforms using PI(4,5)P_2_ as a substrate (Whitman et al., 1988; Traynor-Kaplan et al., 1988; Whitman et al., 1985). PI(3,4,5)P_3_ is transient and its signaling is turned off by the 3’-phosphatase PTEN or the 5’-phosphatase SHIP2 (Maehama and Dixon, 1998; Pesesse et al., 1998). While most PI(3,4,5)P_3_ signaling is associated with the plasma membrane, a significant amount is found on internal vesicles including the nuclear envelope and early endosomes for localized Akt activation (Jethwa et al., 2015). PI(3,4,5)P_3_ is also found on recycling endosomes for AP-1B dependent sorting (Fields et al., 2010), lysosomes to activate mTORC1 (Hsu et al., 2000), and trans-Golgi vesicles where PI(3,4,5)P_3_ on VLDL (very low density lipoprotein) binds to the cargo receptor Sortilin/Vps10 (Sparks et al., 2016, 2020). At the plasma membrane itself, PI(3,4,5)P_3_ recruits the protein kinases Akt and PDK1 to the membrane by engaging their PH domains (Dieterle et al., 2014; Levina et al., 2022).

By homology, *Saccharomyces cerevisiae* lacks p110, thus PI(3,4,5)P_3_ is thought not to exist. However, the fission yeast *Schizosaccharomyces pombe* also lacks a p110 ortholog, yet its class-III Vps34 ortholog can make PI(3,4,5)P_3_ from PI(4,5)P_2_ in addition to its canonical product PI3P (Mitra et al., 2004). This suggests that a synthesis pathway for PI(3,4,5)P_3_ could have evolved prior to the rise of class-I PI 3-kinases as suggested by Mitra and colleagues. Using *S. cerevisiae* vacuoles we examined a role for PI(3,4,5)P_3_ during vacuole fusion. This study shows that PI(3,4,5)P_3_ is made on vacuoles through a Vps34-dependent pathway and is required for vacuole fusion. Removing PI(3,4,5)P_3_ with PTEN or blocking it with the Grp1-PH domain inhibited Ypt7-mediated vertex domain assembly leading to reduced trans-SNARE pairing and fusion. Finally, we found that Vam7 binds PI(3,4,5)P_3_ and is released from membranes when the lipid is modified by PTEN or blocked by Grp1-PH.

## Results

### Short chain PI(3,4,5)P_3_ inhibits in vitro vacuole fusion

In previous studies we have used dioctanoyl (C8) lipids to compete with endogenous sources of their long chain counterparts for binding vacuolar proteins. This revealed that C8-PA competes for Sec18 binding during pre-priming (Starr et al., 2016, 2019; Sparks et al., 2019), and that C8-PI(3,5)P_2_ competes for binding to the V-ATPase subunit Vph1 and regulatory factors of Ca^2+^ transport in a phase between SNARE pairing and hemifusion (Miner et al., 2019, 2020; Zhang et al., 2022). In this study we asked whether adding C8-PI(3,4,5)P_3_ affected vacuole fusion. While this lipid has not been detected in *Saccharomyces cerevisiae*, numerous screening papers have shown that baker’s yeast proteins can bind PI(3,4,5)P_3_, including Vam7 (Yu and Lemmon, 2001; Gallego et al., 2010; Zhu et al., 2001; Dunn et al., 2004). This could be due to lack of specificity in these assays, or it could indicate that the lipid may exist in small transient pools that have escaped detection.

Here we added exogenous C8-PI(3,4,5)P_3_ to in vitro vacuole fusion reactions and found that it potently inhibited fusion with an IC_50_ ∼80 µM **(Fig. 1A).** This showed that vacuole fusion was more sensitive to C8-PI(3,4,5)P_3_ compared to C8-PI(3,5)P_2_ with an IC_50_ ∼140 µM or C8-PI3P, which failed to fully inhibit fusion at 500 µM (Miner et al., 2019). This is an underestimate as C8 lipids only transiently associate with membranes. C8-PIs are mostly aqueous but have a low transient rate of insertion into membranes. Collins and Gordon (2013) calculated that 340 µM C8-PI(4,5)P_2_ was needed to reach ∼4% incorporation (Collins and Gordon, 2013). Thus, C8-PI(3,4,5)P_3_ at its IC_50_ of 80 µM meant that ∼0.9% (0.8 µM) associated with vacuoles at steady state.

**Figure 1.**
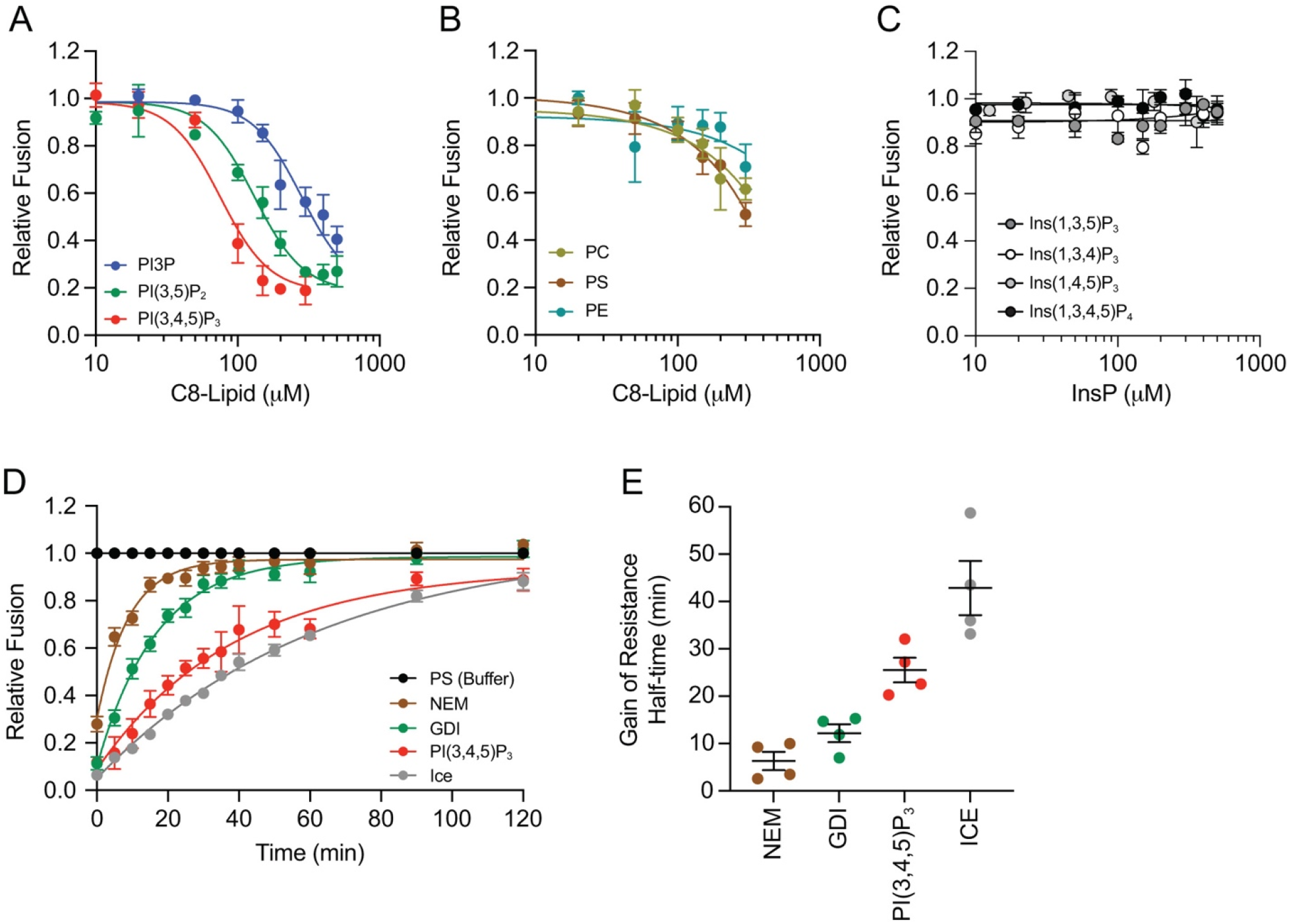
C8-PI(3,4,5)P_3_ inhibits fusion after vacuole tethering. Fusion reactions were treated with buffer alone or curves of: C8-PI3P, C8-PI(3,5)P_2_ and C8-PI(3,4,5)P_3_ **(A);** C8-PC, C8-PS and C8-PE **(B);** Ins(1,3,5)P_3_, Ins(1,3,4)P_3_, Ins(1,4,5)P_3_ and Ins(1,3,4,5)P_4_ **(C)** and incubated for 90 min at 27°C. Fusion was normalized to the maximum fusion (buffer alone) set to 1 for each curve. Data show the mean and SE (n≥3) and fit to one-phase decay curves. IC_50_ values were determined using GraphPad Prism-10. **(D)** Gain of resistance fusion reactions were performed with PS buffer, 1 mM NEM, 2 µM GDI or 150 µM C8-PI(3,4,5)P_3_. Individual reactions were treated with reagents/buffer at the indicated time points. A second set of buffer-treated reactions were placed on ice at the indicated times. Reactions were incubated for a total of 120 min at 27°C. Fusion levels for each reaction were normalized to the untreated control at each time point set to 1. Data show the average and SE (n≥3) and fit to one-phase decay curves. **(E)** Calculated half-times of resistance from assays in **(D)**. Error bars represent SEM (n=3).

To confirm that the measured impacts were PI-specific and not an artifact of the C8 chains, we tested the C8 variants of PC, PE and PS and found that none of the bulk lipids had a significant effect on fusion **(Fig. 1B)**, indicating that the C8 chains themselves had no effect and that the PI headgroups were responsible for altering vacuole fusion. While many PI binding domains, such as the Plc1δ PH, insert a loop into the membrane to stabilize their interaction (Herzog et al., 2016; Lemmon, 2008), they can also simply bind headgroups in solution, albeit with different affinities (Kavran et al., 1998; Ferguson et al., 2000). Based on this, we asked if the headgroups alone could interfere with vacuole fusion. We tested the PI(3,4,5)P_3_ head group Ins(1,3,4,5)P_4_ as well as Ins(1,3,4)P_3_, Ins(1,3,5)P_3_, and Ins(1,4,5)P_3_, and saw that they had no effect at any tested concentration **(Fig. 1C).** This suggested that the head group alone was insufficient for vacuole fusion interference, and that requiring PI(3,4,5)P_3_ likely involves interactions with the membrane phosphate layer.

In order to determine which stage of fusion was affected by C8-PI(3,4,5)P_3_, we performed temporal gain of resistance experiments (Mayer et al., 1996; Haas et al., 1995; Ungermann et al., 1998b; Price et al., 2000b; Sasser et al., 2012; Miner et al., 2019). Inhibitors were added at different timepoints during the assay and incubated for a total of 120 min. As reactions passed a stage of fusion (e.g., Sec18-mediated priming) they became resistant to inhibitors of that stage, such as antibodies against Sec18 and Sec17, NEM, propranolol, or the small molecule IPA (inhibitor of priming activity) (Mayer et al., 1996; Sasser et al., 2012; Starr et al., 2016; Sparks et al., 2019). In this study we used NEM to inhibit Sec18 function and mark the priming threshold, and GDI to inhibit Ypt7 function and mark the tethering/docking phase (Mayer and Wickner, 1997; Haas et al., 1995). Tethering and docking cannot be separated by this assay. Control reactions were treated with buffer alone at 27°C for the duration of the experiment, while a group of buffer-treated tubes were removed and placed on ice to mark the maximum amount of fusion for any recorded time point. We added 150 µM C8-PI(3,4,5)P_3_ to individual reactions at the indicated times to see which stage would be impacted. We found that the gain of resistance curve for C8-PI(3,4,5)P_3_ was shifted to the right of GDI, indicating that it continued to affect fusion after Ypt7-mediated tethering **(Fig. 1D)**. This was further illustrated when the half-times of resistance were calculated. **Fig.1E** shows that the half-times of priming and docking were 5 and 12 min, respectively, whereas the half-times of C8-PI(3,4,5)P_3_ was ∼26 min. This half-time was later than the T_1/2_ ∼15 min we reported for PI(3,5)P_2_ (Miner et al., 2019). It is also significantly later than the half-time of binding PI3P with the Hrs FYVE domain or the Vam7 PX domain alone, which overlaps with the docking curve (GDI or anti-Vam3) as shown by the Ungermann group (Boeddinghaus et al., 2002). Importantly, it should be noted that gain of resistance assays only marks the last step when a variable molecule had an effect, implying earlier stages could have been impacted as well.

### PI(3,4,5)P_3_-specific inhibitors block vacuole fusion

The data presented above suggested that a key PI(3,4,5)P_3_-protein interaction needed for fusion could have been disrupted by exogenous C8-PI(3,4,5)P_3_. To further test for a required PI(3,4,5)P_3_-protein interaction, we used purified GST-Grp1-PH, a PI(3,4,5)P_3_-binding PH domain from the ARF GEF Grp1 (Guillou et al., 2007; Chen et al., 2012; Corbin et al., 2004; Dowler et al., 2000). Sequestering PI(3,4,5)P_3_ from natural binding partners with GST-Grp1-PH inhibited fusion with an average IC_50_ ∼1 µM. This further indicated that the lipid was present on vacuolar membranes **(Fig. 2A)**. We also tested the Grp1-PH^K273A^ mutant that has reduced PI(3,4,5)P_3_ binding affinity (Lindsay et al., 2006; Naughton et al., 2016; Guillou et al., 2007; Yamamoto et al., 2020). We found that it only partially affected fusion. To verify how well Grp1-PH and Grp1-PH^K273A^ bound to PI(3,4,5)P_3_ we performed Biolayer Interferometry (BLI) experiments. For this we used biotinylated PI(3,4,5)P_3_ (b-PI(3,4,5)P_3_) and b-PI3P bound to streptavidin biosensors, which were incubated with 100, 200, 400 and 800 nM Grp1-PH or Grp1-PH^K273A^. We show that Grp1-PH bound to b-PI(3,4,5)P_3_ with a *K_d_* of ∼180 nM whereas b-PI3P was not bound. The Grp1-PH^K273A^ failed to bind either lipid **(Fig. 2B)**. This further suggested that fusion inhibition by Grp1-PH was due to binding PI(3,4,5)P_3_ while the minor effects of Grp1-PH^K273A^ were non-specific.

**Figure 2.**
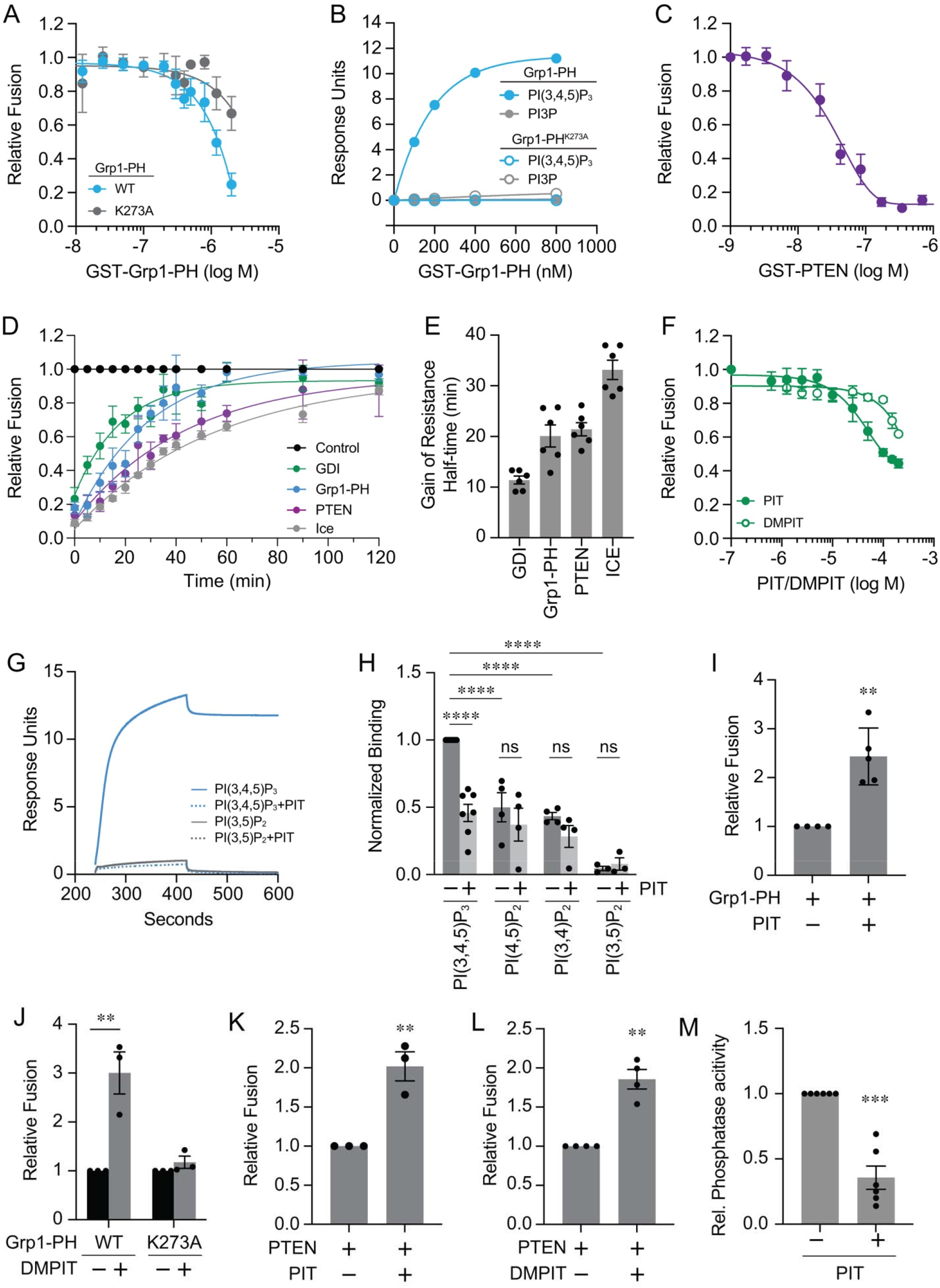
PIP3 specific reagents inhibit vacuole fusion. **(A)** Fusion reactions were treated with buffer or curves of GST-Grp1-PH or GST-Grp1-PH^K273A^ and incubated for 90 min at 27°C. Each curve was normalized to the maximum fusion (no treatment) set to 1. Data shows the averages and SE (n≥3). Data sets were fit to a one-phase decay curve, and IC_50_ values were determined using GraphPad Prism-10. **(B)** BLI of GST-Grp1-PH and GST-Grp1-PH^K273A^ binding to b-PI(3,4,5)P_3_ or b-PI3P. **(C)** Dose curve of GST-PTEN inhibition of fusion as described in (A). **(D)** Gain of resistance fusion reactions were performed with buffer, 1 mM NEM, 2 µM GDI, 2 µM GST-Grp1-PH, or 200 nM PTEN. Reagents/buffer were added at the indicated times. A second set of untreated reactions was placed on ice at each timepoint. Fusion reactions were incubated for a total of 120 min and fusion for each reaction was normalized to the untreated control for the each timepoint at 27°C set to 1. Data sets show the average and SE and fit to one-phase decay curves (n≥3). **(E)** Calculated half-times of resistance from assays in **(D)**. **(F)** Dose curves of PIT-1 and 3,5-dimethyl PIT-1 (DMPIT) on vacuole fusion. **(G)** Sensorgrams of BLI and PIT-1 (200 µM) inhibition of Grp1-PH binding to b-PI(3,4,5)P_3_ and b-PI(3,5)P_2_. **(H)** Average Grp1-PH binding to b-PI(3,4,5)P_3_, b-PI(4,5)P_2_, b-PI(3,4)P_2_, and b-PI(3,5)P_2_ and the effects of 200 µM PIT-1. Data was normalized to Grp1-PH binding to b-PI(3,4,5)P_3_ set to 1. Error bars represent mean ± SE. (n≥3). Significance was determined by unpaired one-way ANOVA for multiple comparisons. Tuckey’s post-hoc test was used to determine individual *p*-values. \*\*\*\**p*<0.0001. ns, not significant. **(I-M)** PIT-1 (200 µM) and DMPIT (200 µM) reversal of fusion inhibition by 2.3 µM WT Grp1-PH or Grp1-PH^K273A^ **(I-J)** and 200 nM PTEN **(K-L)**. Fusion efficiency was normalized to Grp1-PH or PTEN set to 1. Error bars represent mean ± SE (n≥3). **(I-M)** Significance was determined unpaired two-tailed t-test. \*\**p*<0.01. **(M)** Fluorescent phosphatase assay of PTEN and C8-PI(3,4,5)P3 in the presence or absence of 200 µM PIT-1. Error bars represent mean ± SE (n≥3). Significance was determined unpaired two-tailed t-test. \*\*\*\**p*<0.0001.

To further interrogate the function of PI(3,4,5)P_3_ on vacuoles we used the lipid phosphatase PTEN, which converts PI(3,4,5)P_3_ to PI(4,5)P_2_ (Maehama and Dixon, 1998; Lee et al., 1999). We added a dose-response curve of GST-PTEN to fusion reactions and found that it inhibited with an IC_50_ ∼60 nM **(Fig. 2C)**. This further signified that endogenous vacuolar PI(3,4,5)P_3_ affected fusion. While PTEN does have some activity against PI(3,4)P_2_ and PI3P, it is far weaker than its activity against PI(3,4,5)P_3_ (Lee et al., 1999).

We then tested if PTEN and Grp1-PH had gain of resistance curves that matched the C8-PI(3,4,5)P_3_ curve. Using the aforementioned assay, we found that the resistance curves of Grp1-PH and PTEN had similar half-times with means of ∼20-22 min, which were consistent with the C8-PI(3,4,5)P_3_ data **(Fig. 2D-E)**. Interestingly, after their resistance half-times, the Grp1-PH and PTEN curves diverged so that binding alone by Grp1-PH lost its effect before PTEN, indicating that hydrolysis remained effective for a longer period. Roughly, the T=80% for Grp1-PH and PTEN were 40 and 75 min, respectively.

Next, we asked if we could protect native PI(3,4,5)P_3_ from PTEN and Grp1-PH with a competitive inhibitor. We used PIT-1, a small molecule that binds to the PI(3,4,5)P_3_ binding pockets of PH domains including those of AKT, PDK1, Grp1 and ARNO (Miao et al., 2010). Notably PIT-1 does not interfere with PI(3,4)P_2_ interactions with TAPP1 or TAPP2, or PI(4,5)P_2_ interactions with PLC. First, we tested PIT-1 alone and found that it partially inhibited fusion at 200 µM, which was near the concentration needed for its inhibition of Akt-PH-PI(3,4,5)P_3_ interactions (Miao et al., 2010) **(Fig. 2F)**. Higher concentrations (>200 µM) were not tested due to solubility issues. We also used the PIT-1 derivative 3,5-dimethyl PIT-1 (DMPIT) and observed little interference with fusion **(Fig. 2F)**. The weak effects on fusion could be due to weaker affinities against native protein-lipid affinities.

### PIT-1 blocks Grp1-PH binding to PI(3,4,5)P**_3_**

To directly test the effect of PIT-1 on protein-lipid interactions, we used BLI and biotinylated PI(3,4,5)P_3_, PI(4,5)P_2_, PI(3,4)P_2_, and PI(3,5)P_2_ bound to streptavidin probes. Lipid-bound probes were dipped into wells containing 400 nM Grp1-PH to measure binding kinetics. In **Fig.2G** we show sensorgrams of Grp1-PH binding to lipids in the presence or absence of PIT-1. We found that PIT-1 reduced Grp1-PH binding to b-PI(3,4,5)P_3_ **(Fig. 2H)**. We also found that Grp1-PH interacted with b-PI(4,5)P_2_ and b-PI(3,4)P_2_, albeit at lower levels that were equal to PIT-1-reduced binding of b-PI(3,4,5)P_3_. In addition, we found that Grp1-PH failed to bind b-PI(3,5)P_2_. Importantly, PIT-1 had no effect on the interactions between Grp1-PH and b-PI(4,5)P_2_ or b-PI(3,4)P_2_. This was consistent with its reported specificity for PI(3,4,5)P_3_ binding sites. Interactions with non-specific anionic lipids likely reflected electrostatic binding of basic residues K279 and R284 that are adjacent to the PI(3,4,5)P_3_ binding pocket (Lumb and Sansom, 2012; Lai et al., 2013). These interactions guide and orient Grp1-PH to facilitate finding PI(3,4,5)P_3_ and increase affinity (Corbin et al., 2004).

The lack of interactions of Grp1-PH with b-PI3P and b-PI(3,5)P_2_ could be due to multiple factors. First, it may be that the D4-phosphate was essential for moderate interactions with Grp1-PH due to clustering negative charges with the D3 or D5 phosphates to increase H-bonding and binding energy (Sohn et al., 2018; Li et al., 2013). On the other hand, the separation of phosphates in PI(3,5)P_2_ could have changed the number of contacts with Grp1-PH to prevent binding. This is underscored by their distinct interactomes (Catimel et al., 2008). The lack of binding to b-PI3P was most likely due to insufficient contact points of the monophosphate with Grp1. PI5P and PI4P were not tested.

### PIT-1 blocks fusion inhibition by Grp1-PH and PTEN

Next, we tested if PIT-1 could rescue the inhibition of fusion by Grp1-PH and PTEN. For this, we pre-mixed Grp1-PH and PIT-1 for 5 min at 27°C. Next, fusion reactions were supplemented with Grp1-PH alone or a mixture of Grp1-PH and PIT and incubated for a total of 90 min. This showed that PIT was able to partially block the inhibitory effect of Grp1-PH **(Fig. 2I)**. Likewise, we found that DMPIT partially blocked the effect of Grp1-PH **(Fig. 2J)**. To see if this was due to PI(3,4,5)P_3_, we asked if the minor effects of the Grp1-PH^K273A^ on fusion would be affected by DMPIT. We show that DMPIT had no effect on Grp1-PH^K273A^, further demonstrating that the effects of Grp1-PH and PIT-1/DMPIT required PI(3,4,5)P_3_. This further indicated that the effects of Grp1-PH^K273A^ on fusion were independent of PI(3,4,5)P_3_.

Finally, we asked if PIT-1 or DMPIT could reverse the inhibitory effects of PTEN on fusion. Using the same approach, we pre-mixed PTEN and either PIT-1 or DMPIT for 5 min at 27°C. Next, fusion samples received PTEN alone or the mixture and were incubated for 90 min at 27°C. This showed that PIT-1 and DMPIT could partially block the inhibitory effects of PTEN **(Fig. 2K-L)**. While PTEN does not have a PH domain, it must interact with PI(3,4,5)P_3_, indicating that PIT-1 is not specific for PH domains, and that other PI(3,4,5)P_3_ interaction sites could be inhibited. To verify if the block of PTEN fusion inhibition by PIT was through altering phosphatase activity we used a fluorescent phosphatase assay. This was used in lieu of the standard malachite green assay due to the interference of PIT-1 with the colorimetric assay. We found that incubating PTEN with C8-PI(3,4,5)P_3_ and the fluorescent phosphate reporter caused a sharp increase in fluorescence, indicating that PTEN phosphatase activity released a Pi from the lipid **(Fig. 2M)**. When PIT-1 was introduced to the reaction, we found that the signal was decreased, demonstrating that PTEN phosphatase activity was inhibited. These experiments only used pure components in the absence of vacuoles, indicating that PIT-1 directly blocked C8-PI(3,4,5)P_3_ hydrolysis by PTEN.

### PI(3,4,5)P_3_ is present on isolated vacuoles

The inhibition of fusion by PTEN and Grp1-PH signified that PI(3,4,5)P_3_ was present on vacuoles. To visualize where PI(3,4,5)P_3_ was localized on docked vacuoles, we used subinhibitory concentrations of GST-Grp1-PH (150 nM) and fluorescent (CF488) anti-GST antibody (CF488-Grp1-PH). This prevented any interference caused by conjugating primary amines or free Cys with reactive dyes. Grp1-PH has key Lysines in the lipid binding pocket and a Cysteine next to a lipid-interacting Lys (Cronin et al., 2004; Lai et al., 2013). Analysis of the relative distribution of CF488-Grp1-PH was executed as previously described (Wang et al., 2002; Eitzen et al., 2002; Wang et al., 2003; Fratti et al., 2004; Karunakaran et al., 2012; Jun et al., 2006; Miner et al., 2019, 2020; Zhang et al., 2024). Vacuoles were stained with the vital dye FM4-64 to label the entirety of membranes (Magenta) and CF488-Grp1-PH to label PI(3,4,5)P_3_ (Green). Fluorescence microscopy showed that CF488-Grp1-PH was enriched at the vertex microdomains of docked vacuoles where essential lipids and proteins accumulate to trigger fusion (Starr and Fratti, 2019) **(Fig. 3A).** Arrows marked examples of enriched vertex sites whereas arrow heads pointed at outer edge representatives. A heat map showed fluorescence intensities of CF488-Grp1-PH at individual vertex sites. Quantitation of vertex enrichment was determined by taking the ratios of CF488-Grp1-PH to FM4-64 fluorescence intensities at vertices versus outer edge membranes not in contact with other vacuoles. These were taken for each vertex in a cluster. Ten-to-fifteen clusters were measured for each condition per experiment for a total of three experiments. The outer edge ratios were normalized to 1 and the enrichment of CF488-Grp1-PH at vertices were expressed relative to outer edge intensities. The vertex and outer membrane ratios of CF488-Grp1-PH:FM4-64 (specific/non-specific) were charted in a cumulative distribution graph where individual measurements were ranked and plotted at their rank percentile **(Fig. 3B)**. The geometric means and geometric standard deviations were also determined for statistical analysis **(Fig. 3C)**. This showed that CF488-Grp1-PH was indeed enriched at vertices relative to outer edges.

**Figure 3.**
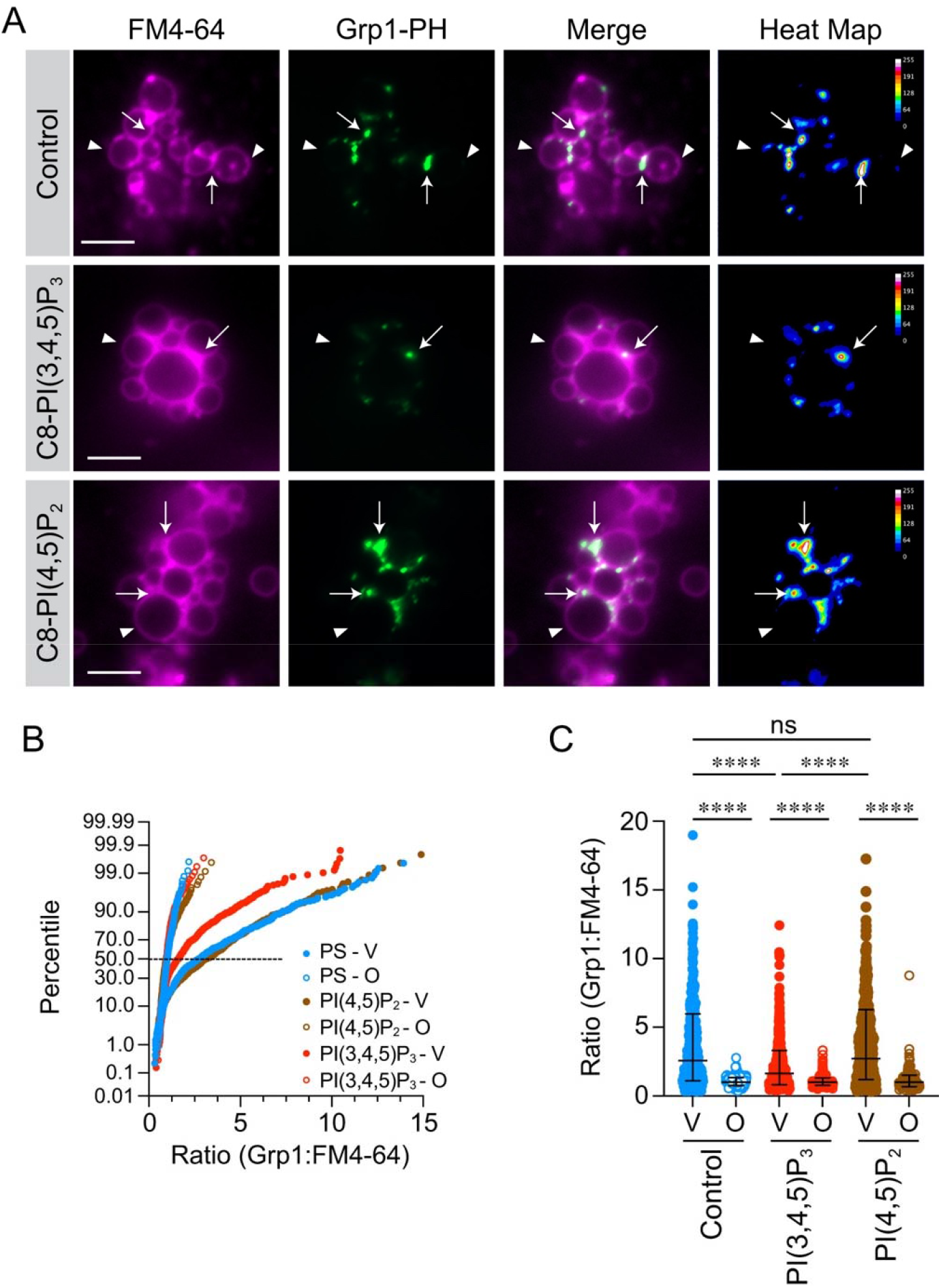
PI(3,4,5)P_3_ competes for Grp1-PH binding. **(A)** Docking reactions were performed with 300 nM C8-PI(4,5)_2_, 300 nM C8-PI(3,4,5)P_3_, or buffer to compete for GST-Grp1-PH labeling. Membranes were labeled with FM4-64 (magenta), and 150 nM GST-Grp1-PH. GST was visualized with CF488-anti-GST IgG (Green). Vacuoles were incubated at 27°C for 30 min then processed for imaging as described in the methods section. Channels were merged to show vertex enrichment (Merge). Heat maps were generated in ImageJ to show CF488 relative intensities with a calibration bar. **(B)** Data points were pooled from multiple experiments and each experiment contained ratios from 10 or more fields with ≥10 clusters made of ≥5 vacuoles. Data was ordered in cumulative distribution plots to depict the percentile of specific/non-specific ratio for vertices (V) and outer edge membranes (O) (Control-V = 432; Control-O = 240; PI(3,4,5)P_3_-V = 624; PI(3,4,5)P_3_-O; PI(4,5)P_2_-V = 441; PI(4,5)P_2_-O = 228). **(C)** Quantitation of ratiometric fluorescence intensities of vertices and outer edge in panel B. Data points were pooled from multiple experiments and presented as scatter plots. Error bars represent geometric means ± geometric SD (n=3). Significance was determined using one-way ANOVA for multiple comparisons and Tukey’s test for individual *p*-values. \*\*\*\**p*<0.0001. ns, not significant. Scale bars: 5 µm.

To verify that CF488-Grp1-PH staining was due to PI(3,4,5)P_3_ binding we performed competition experiments. For this, we added 300 nM C8-PI(3,4,5)P_3_, or an equal volume of PS buffer as described. This showed that C8-PI(3,4,5)P_3_ significantly reduced CF488-Grp1-PH staining relative to the control, indicating that it competed with the endogenous lipid for binding. This level of C8-PI(3,4,5)P_3_ does not affect fusion and only competed with Grp1-PH for labeling. As a negative control, we used 300 nM C8-PI(4,5)P_2_ and found that it had no effect on CF488-Grp1-PH labeling. This further demonstrated that CF-488-Grp1-PH labeling was PI(3,4,5)P_3_-dependent.

### PTEN and Tep1 affect PI(3,4,5)P_3_ enrichment at vacuoles

Earlier, we showed that PTEN inhibited fusion and that its effects were partially blocked by PIT-1. Here, we tested how PTEN affected PI(3,4,5)P_3_ localization on vertices. We incubated docking reactions with 50 nM PTEN or PS buffer for 30 min at 27°C as described. This showed that PTEN reduced the number and intensity of CF488-Grp1-PH labeled vertices, indicating that PI(3,4,5)P_3_ levels were reduced **(Fig. 4A-C)**. To expand on this, we looked at the role of the yeast PTEN ortholog Tep1. While Tep1 plays a role in sporulation, it is considered to be inactive as a lipid phosphatase and unimportant for normal growth (Heymont et al., 2000). This was based on the lack of a growth defect when deleted in haploid cells. The growth of the homozygous diploid deletion was resistant to the pan PI3K inhibitors wortmannin and LY294002; however, PI(3,4,5)P_3_ and PI(4,5)P_2_ levels were not examined. Nevertheless, we next asked if vacuoles isolated from the haploid *tep1*Δ cells had altered CF488-Grp1-PH labeling. The results showed that deleting *TEP1* led to increased CF488-Grp1-PH labeling relative to the WT parent, indicating that the loss of Tep1 caused a significant accumulation of PI(3,4,5)P_3_ **(Fig. 4D-F)**. The increase in PI(3,4,5)P_3_ in *tep1*Δ vacuoles complemented the reduction in lipid when PTEN was added.

**Figure 4.**
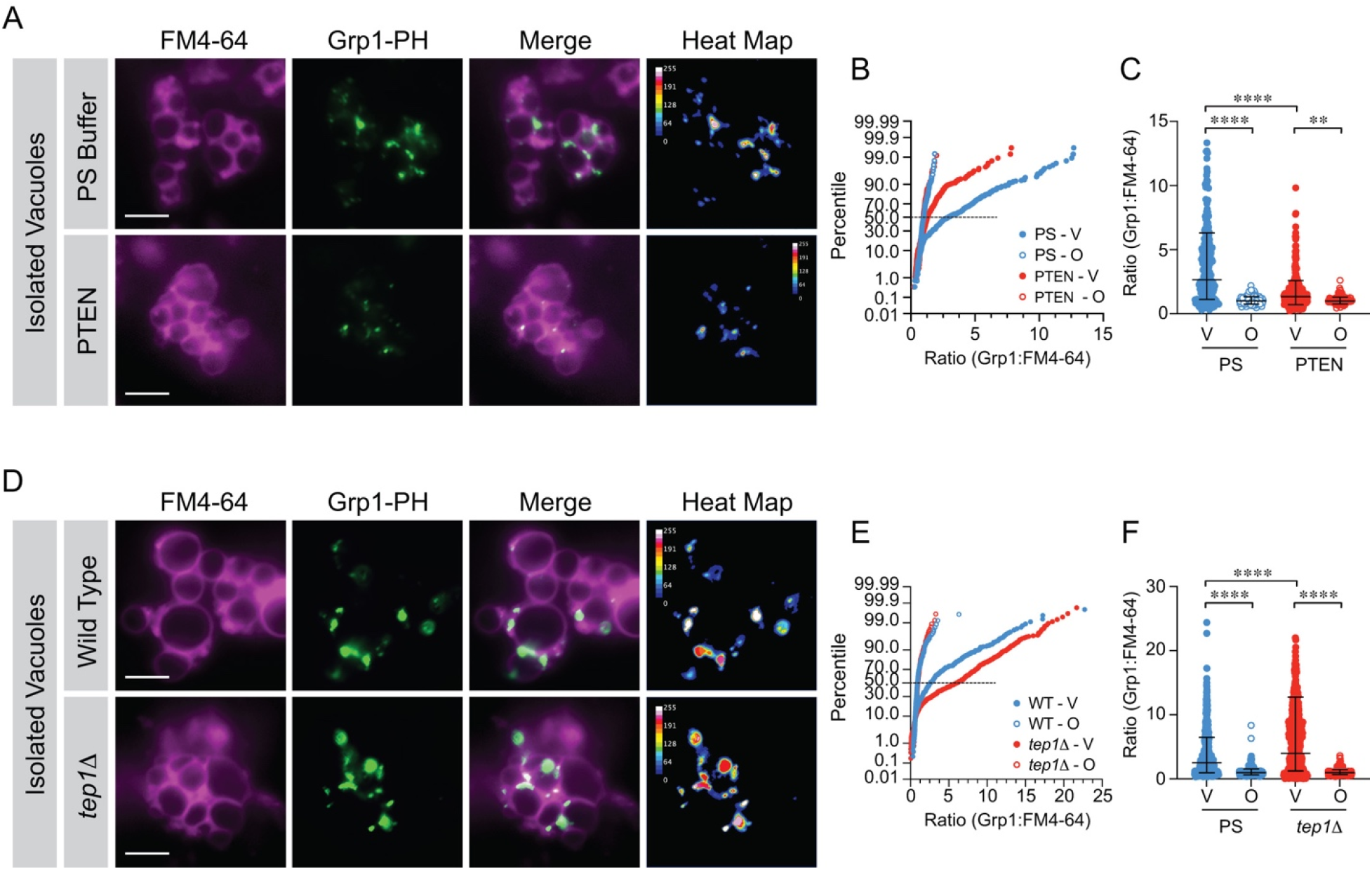
PTEN reduces Grp1-PH vacuole staining whereas and *TEP1* deletion augments staining. **(A)** Docking reactions were treated with 50 nM PTEN or PS buffer (Control) and incubated for 30 min at 27°C. At the end of the incubation period, reactions were placed on ice, labeled with FM4-64 and prepared for fluorescence microscopy as described in Fig.3. **(B)** Data points were pooled from three experiments and ordered by cumulative distribution plots to depict as described in Fig.3 (Control-V = 275; Control-O = 142; PTEN-V = 275; PTEN-O = 121). **(C)** Quantitation of ratiometric fluorescence intensities of vertices and outer edge in panel B. Data points were pooled from multiple experiments and presented in a scatter plot. **(D)** Docking reactions of WT vacuoles and *tep1*Δ vacuoles were incubated for 30 min at 27°C then processed for imaging as described above. **(E)** Cumulative distribution plots of pooled data from three experiments (WT-V = 447; WT-O = 255; *tep1*Δ-V = 559; *tep1*Δ-O = 263). (F) Scatter plots of pooled data from **(E)**. Error bars represent geometric means ± geometric SD. Significance was determined using one-way ANOVA for multiple comparisons and Tukey’s test for individual *p*-values. \*\**p*<0.01, \*\*\*\**p*<0.0001. Scale bars: 5 µm.

### Vps34 is required for PI(3,4,5)P_3_ production

While *S. cerevisiae* lacks a class I PI3K p110 ortholog, there is a precedent for PI(3,4,5)P_3_ production by the class III PI3K Vps34 ortholog in *S. pombe* (Mitra et al., 2004). To examine if Vps34 was involved in PI(3,4,5)P_3_ production, we use the class-specific inhibitor SAR405 (Ronan et al., 2014). Vacuole labeling with CF488-Grp1-PH was significantly reduced on vacuoles treated with SAR405 **(Fig. 5A-C)**, which signified that PI(3,4,5)P_3_ production occurred in vitro on isolated vacuoles, in a Vps34-dependent manner, and not due to an unidentified class-I PI3K ortholog.

**Figure 5.**
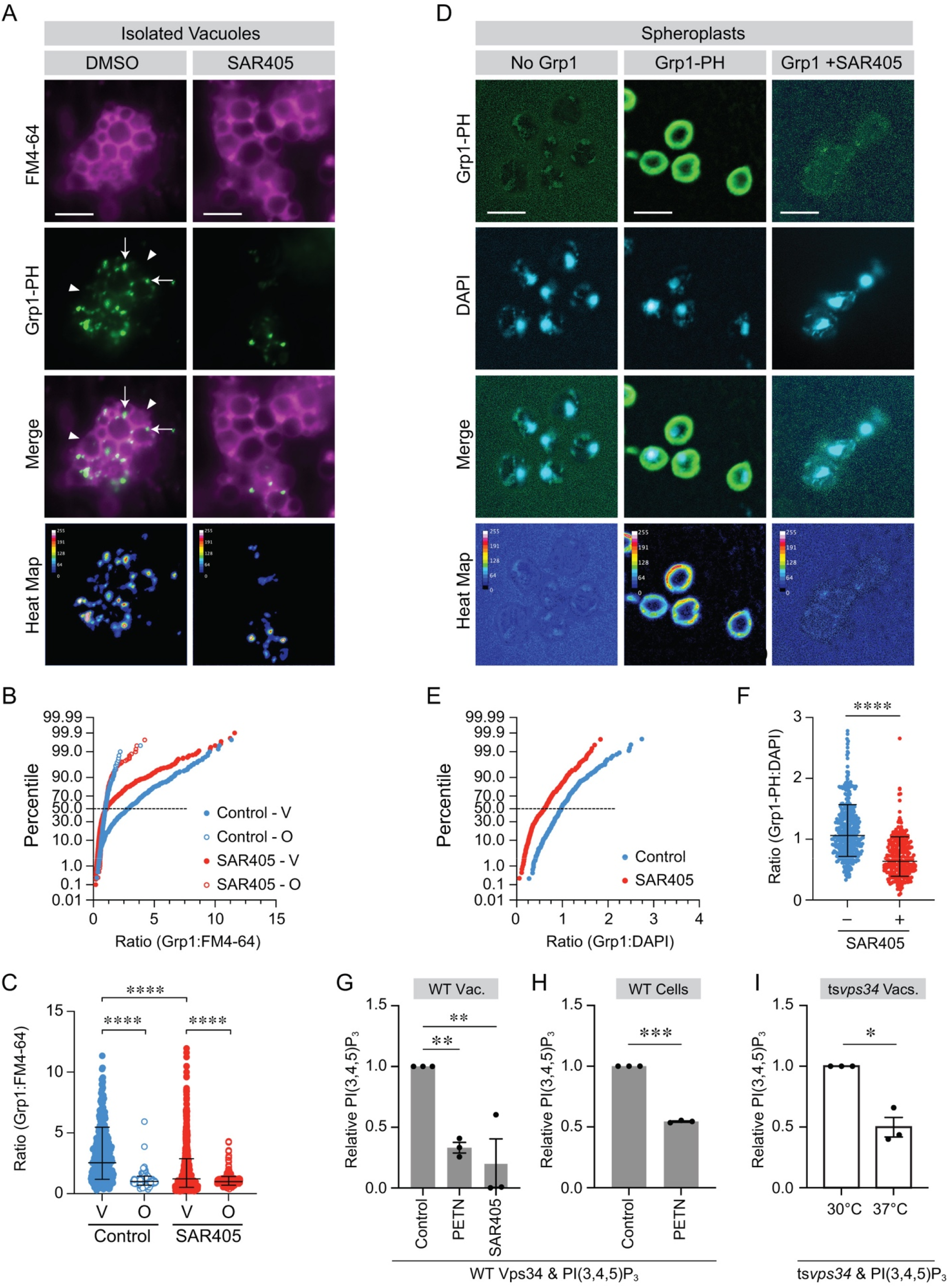
PI(3,4,5)P_3_ production requires Vps34 on isolated vacuoles and the plasma membrane. **(A)** Docking reactions of purified vacuoles were incubated for 30 min at 27°C with and labeled with GST-Grp1-PH as described in Fig.3. Reactions were treated with 200 µM SAR405 or DMSO and prepared for fluorescence microscopy. After incubation the reactions were placed on ice and mixed with 5 µM FM4-64 to label entire vacuoles. Vacuoles were pelleted to remove excess antibody and resuspended in PS buffer. Reactions were next mixed with low melt agarose, mounted on glass slides and observed by fluorescence microcopy. Scale bars: 5 µm. **(B)** Cumulative distribution plot of pooled experiments (n=3) (DMSO-V = 403; DMSO-O = 200; SAR405-V = 990; SAR405-O = 394). **(C)** Pooled data shown as a scatter plot with geometric means and their geometric standard deviations. Significance was determined using one-way ANOVA for multiple comparisons and Tukey’s test for individual *p*-values. \*\*\*\**p*<0.0001. **(D)** Mid-log cells were fixed with formaldehyde and spheroplasted to detect PI(3,4,5)P_3_. Spheroplasts were incubated with 10 µM GST-Grp1-PH or buffer followed by CF488 anti-GST antibody. Reactions were treated with 200 µM SAR405 or DMSO and incubated for at 30°C. Samples were next mounted on slides with Fluoromount-G containing DAPI to label nuclei. Separate samples were incubated with anti-GST in the absence of Grp1-PH. GST is shown in Green and DAPI is shown in Blue. Heat maps show GST-Grp1-PH enrichment over background. Scale bars: 5 µm. **(E)** Cumulative distribution plots of Grp1:DAPI fluorescence ratios of cells treated with SAR405 (n=413) or DMSO (n=465). **(F)** Quantitation of plasma membrane labeling with CF488-Grp1-PH. Ratios of maximum fluorescence intensities (CF488:DAPI) were determined for ≥100 cells per condition per experiment (n=4). Significance was determined using unpaired two-tailed t-test. \*\*\*\**p*<0.0001. **(G)** Quantitation of vacuole PI(3,4,5)P_3_ was performed using an ELISA kit. Fusion reactions were treated with 150 nM PTEN or 200 µM SAR405 and incubated for 30 min at 27°C. **(H)** ELISA quantitation of WT cell PI(3,4,5)P_3_ in the presence or absence of PTEN. **(I)** ELISA quantitation of ts*vps34* vacuole PI(3,4,5)_3_ at 30°C or 37°C. Bars represent mean ± SE (n=3). Significance was determined by unpaired two-tailed t-test or one way ANOVA for multiple comparisons and Tukey’s test for individual *p*-values. \**p*<0.05. \*\**p*<0.01, \*\*\**p*<0.001.

Although SAR405 reduced the PI(3,4,5)P_3_ signal on isolated vacuoles, a significant number of vertices stained with CF488-Grp1-PH, indicating that either pre-existing lipid was present on membranes or that there was an incomplete shutdown of activity. Pre-existing lipid could come from vacuolar production prior to isolation as well as trafficking from the plasmalemma (Sahan et al., 2025). To test if the plasma membrane contained PI(3,4,5)P_3_ we used whole cells that were treated with lyticase to digest the cell wall and permeabilize the plasma membrane, and fixed with formaldehyde (Martínez-Muñoz and Peña, 2005; Pringle et al., 1991). Cells were then incubated with CF488-Grp1-PH, washed and mounted with media containing DAPI to label nuclei. **Fig.5D** shows fluorescence for individual channels as well as a merged image and a heat map to show relative CF488-Grp1-PH intensities. Our data showed that CF488-Grp1-PH labeled the plasmalemma, indicating that PI(3,4,5)P_3_ was present. In the absence of Grp1-PH, the anti-GST IgG failed to label the spheroplasts. When cells were pretreated with SAR405, the CF488-Grp1-PH labeling was strongly reduced. CF488-Grp1-PH staining intensities were quantified by ratiometric fluorescence measurements of CF488 and DAPI. This showed that untreated cells had a mean fluorescence that was significantly greater compared to those treated with SAR405 **(Fig. 5E-F)**. This indicated that PI(3,4,5)P_3_ was enriched at the plasmalemma and that it was produced in a Vps34-dependent manner. While PI(3,4,5)P_3_ was on the plasmalemma, this did not necessarily show that it was made on the membrane or whether it was trafficked from other organelles.

Spheroplast staining failed to label vacuolar PI(3,4,5)P_3_, which was likely due to the excess of plasmalemma content that masked the vacuolar signal. This was reminiscent of differential PI(4,5)P_2_ staining between the plasmalemma and vacuoles. Plasma membrane PI(4,5)P_2_ was in great excess, preventing the detection of vacuole staining labeling (Stefan et al., 2002). Vacuolar PI(4,5)P_2_ was only detected when the organelles were isolated from the cell (Mayer et al., 2000; Fratti et al., 2004). PI(4,5)P_2_ was also seen on vacuoles when the phosphatases Inp54 and Sac1 were deleted (Wiradjaja et al., 2007).

### PI(3,4,5)P_3_ quantitation

To compare vacuolar versus whole cell PI(3,4,5)P_3_, we extracted lipids and analyzed their concentration using a PI(3,4,5)P_3_ mass ELISA kit (Choi et al., 2016; Gross et al., 2015). For vacuole measurements, we extracted lipids from 3X docking reactions (18 µg by protein) which contained an average of ∼1 pmol PI(3,4,5)P_3_ per 10^6^ cells **(Fig. 5G)**. Due to the range in daily lipid extraction efficiency, we normalized each experiment to the control set to 1. Treatment with 150 nM PTEN prior to lipid extraction reduced PI(3,4,5)P_3_ levels by >70%. In parallel, vacuoles that were treated with 200 µM SAR405 also had greatly reduced PI(3,4,5)P_3_. Next, we extracted lipids from whole cells grown to 0.65 O.D._600_ and found that they contained ∼31 pmol PI(3,4,5)P_3_/10^6^ cells which decreased to ∼18 pmol/10^6^ cells when treated with PTEN **(Fig. 5H)**. The difference between whole cell and vacuole PI(3,4,5)P_3_ was ∼32-fold in cells (containing vacuoles) compared to vacuoles alone.

### PI(3,4,5)P_3_ production and Vps34

The SAR405 data indicated that a class III PI3K was required for making PI(3,4,5)P_3_, thus it was important to directly test Vps34 as the sole member of the class. To get a better sense of Vps34 activity in PI(3,4,5)P_3_ production, we used cells that expressed a temperature sensitive mutation (ts*vps34*) (Herman and Emr, 1990; Stack et al., 1995). Vacuoles were isolated from ts*vps34* cells grown at 30°C. Isolated ts*vps34* vacuoles were then incubated at the permissive (30°C) or non-permissive temperatures (37°C) to modulate Vps34 activity without the need for chemical inhibitors. Lipids were extracted from ts*vps34* vacuoles and quantified by ELISA. This showed that inactivation of ts*vps34* at 37°C resulted in a 50% reduction in PI(3,4,5)P_3_ (0.24 to 0.098 pmol/million cells) **(Fig. 5I)**.

We next wanted to visualize the effects of ts*vps34* inactivation on PI(3,4,5)P_3_ distribution. For this, we isolated vacuoles from ts*vps34* cells grown at 30°C. These vacuoles were then incubated at the permissive (30°C) or non-permissive temperatures (37°C) to modulate Vps34 activity. Reactions were incubated for 30 min, and CF488-Grp1-PH labeling was carried out as described above. Vacuoles incubated at 30°C showed CF488-Grp1-PH vertex labeling whereas those incubated at 37°C had significantly reduced labeling, indicating that Vps34 was required for PI(3,4,5)P_3_ production during the reaction **(Fig. 6A-C)**. To verify that CF488-Grp1-PH staining itself was not temperature sensitive, we repeated the experiment with vacuoles containing WT Vps34. This showed that CF488-Grp1-PH was not sensitive to changes in temperature as it was equally enriched at vertices under both conditions **(Fig. 6D-F)**.

**Figure 6.**
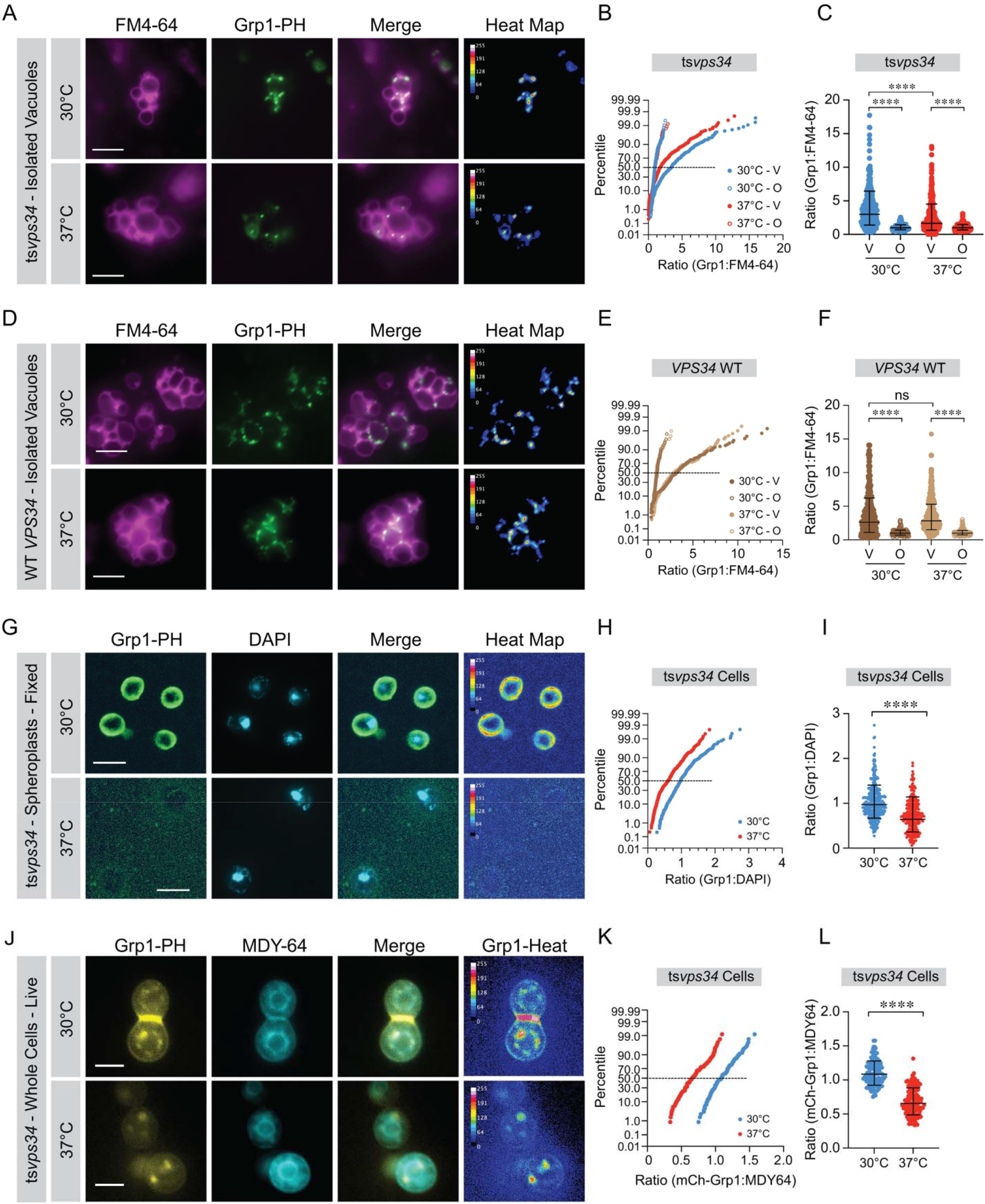
Temperature sensitive ts-vps34. **(A)** Docking reactions using purified ts*vps34* vacuoles were incubated for 30 min at 30°C or 37°C. PI(3,4,5)P_3_ was labeled with GST-Grp1-PH and CF488 anti-GST IgG as described in Fig.3. After incubation the reactions placed on ice and labeled with FM4-64 to show entire vacuoles. Vacuoles were pelleted to remove excess antibody and resuspended in PS buffer. Reactions were next mixed with low melt agarose, mounted on glass slides and observed by fluorescence microcopy. Scale bars: 5 µm. **(B)** Cumulative distribution plot of pooled experiments (n=3) (30°C-V = 328; 30°C-O = 212; 37°C-V = 454; 37°C-O = 260). **(C)** Pooled data shown as a scatter plot with geometric means and their geometric standard deviations. Significance was determined using one-way ANOVA for multiple comparisons and Tukey’s test for individual *p*-values. \*\*\*\**p*<0.0001. **(D)** Docking reactions with WT vacuoles were incubated for 30 min at 30°C or 37°C. and labeled for PI(3,4,5)P_3_ as described above. **(E)** Images were analyzed and plotted in cumulative distribution plots as above (30°C-V = 280; 30°C-O = 116; 37°C-V = 359; 37°C-O = 178). **(F)** Scatter plots of pooled data shown in panel D. Error bars represent geometric means ± geometric standard deviations. Significance was determined using one-way ANOVA for multiple comparisons and Tukey’s test for individual *p*-values. \*\*\*\**p*<0.0001, ns – not significant. **(G)** Mid-log ts*vps34* cells were fixed with formaldehyde and spheroplasted to detect PI(3,4,5)P_3_ as described in Fig.5. Samples were next mounted on slides with Fluoromount-G containing DAPI to label nuclei. GST is shown in Green and DAPI is shown in Blue. Heat maps show GST-Grp1-PH enrichment over background. Scale bars: 5 µm. **(H)** Ratios of maximum fluorescence intensities (CF488:DAPI) were determined for ≥100 cells per condition per experiment (n=3). Cumulative distribution plots of Grp1:DAPI fluorescence ratios of cells treated with 30°C (n=448) or 37°C (n=429). **(I)** Scatter plot of plasma membrane labeling with CF488-Grp1-PH. Significance was determined using unpaired two-tailed t-test. \*\*\*\**p*<0.0001. **(J)** In vivo expression of mCherry-Grp1-PH in ts*vps34* cells. Cells were grown at 30°C or 37°C and stained with MDY-64 to label vacuoles. mCherry-Grp1-PH and MDY-64 were pseudocolored yellow and cyan, respectively. Merged images were created in Photoshop to show colocalization of mCherry-Grp1-PH and MDY-64. Heat maps and calibration bars of mCherry-Grp1-PH fluorescence intensities were created using ImageJ. **(K)** Ratios of maximum fluorescence intensities (mCherry:MDY-64) were determined for ≥50 cells per condition per experiment (n=3). Cumulative distribution plots of cells grown at 30°C (n=153) and 37°C (n=165). **(L)** Scatter plot of plasma membrane labeling with mCherry-Grp1-PH. Significance was determined using unpaired two-tailed t-test. \*\*\*\**p*<0.0001.

To see if Vps34 was also responsible for CF488-Grp1-PH staining at the plasma membrane, we generated spheroplasts as described in the methods section. Spheroplasts were incubated for 1 h in the presence of ATP at 30°C or 37°C. Spheroplasts were fixed with formaldehyde prior to staining with CF488-Grp1-PH and DAPI. We found that inhibiting ts*vps34* at 37°C reduced plasmalemma labeling with CF488-Grp1-PH, while labeling was preserved in cells grown at the permissive temperature **(Fig. 6G-I)**. This indicated that Vps34 was required for PI(3,4,5)P_3_ accumulation at the plasma membrane.

### In vivo PI(3,4,5)P_3_ labeling

In vitro labeling with CF488-Grp1-PH clearly showed that PI(3,4,5)P_3_ was present on isolated vacuoles as well as spheroplast plasmalemma. To see PI(3,4,5)P_3_ in live cells, we constructed a p405ADH1 plasmid containing GST-mCherry-Grp1-PH (mCh-Grp1) under expression control of the constitutive *ADH1* (alcohol dehydrogenase 1) promoter. We used ts*vps34* cells transformed with mCh-Grp1 to localize PI(3,4,5)P_3_. We found that ts*vps34* cells grown at 30°C showed distinct vacuole mCh-Grp1 staining that overlapped with MDY-64 **(Fig. 6J-L)**. We also observed plasma membrane labeling with mCh-Grp1. On the other hand, incubating at 37°C reduced vacuole and plasma membrane mCh-Grp1 labeling. Each fluorescence channel is shown as well as a merged image and a heat map for mCh-Grp1 intensity.

### PI(4,5)P_2_ is the Vps34 substrate for PI(3,4,5)P_3_ production

The data thus far has indicated that Vps34 was required for PI(3,4,5)P_3_ production suggesting that PI(4,5)P_2_ was the substrate. PI(4,5)P_2_ is produced on isolated vacuoles and localizes to vertex microdomains (Mayer et al., 2000; Fratti et al., 2004; Karunakaran et al., 2012). To test if PI(4,5)P_2_ was directly linked to PI(3,4,5)P_3_ production, we used a temperature sensitive mutation of the PI4P 5-kinase Mss4 (ts*mss4*) (Stefan et al., 2002; Audhya and Emr, 2003). First, we tested ts*mss4* vacuoles for PI(4,5)P_2_ production. Vacuoles were isolated from ts*mss4* cells grown at 30°C then split for experiments where vacuoles were incubated under docking conditions for 30 min at 30°C or 37°C. The presence of PI(4,5)P_2_ was detected with the ENTH domain from Epsin1 (Leitner et al., 2019; Hom et al., 2007; Fratti et al., 2004). At the permissive temperature, we observed enriched labeling of vertices with 150 nM recombinant GST-ENTH and CF488 anti-GST IgG (CF488-ENTH) as performed for GST-Grp1-PH labeling **(Fig. 7A-C)**. This level of GST-ENTH had no effect on vacuole fusion as the IC_50_ for blocking fusion was 560 nM **(Supplemental Fig.1D)**. When ts*mss4* vacuoles were incubated at 37°C we observed a sharp decrease in CF488-ENTH labeling, indicating that PI(4,5)P_2_ production was indeed blocked. To control for temperature sensitive ENTH binding, we used WT vacuoles (*MSS4*) incubated at 30°C or 37°C. Similar to WT Vps34 labeling with CF488-Grp1-PH, the shift in temperature had no effect on CF488-ENTH labeling **(Supplemental Fig.1A-C)**.

**Figure 7.**
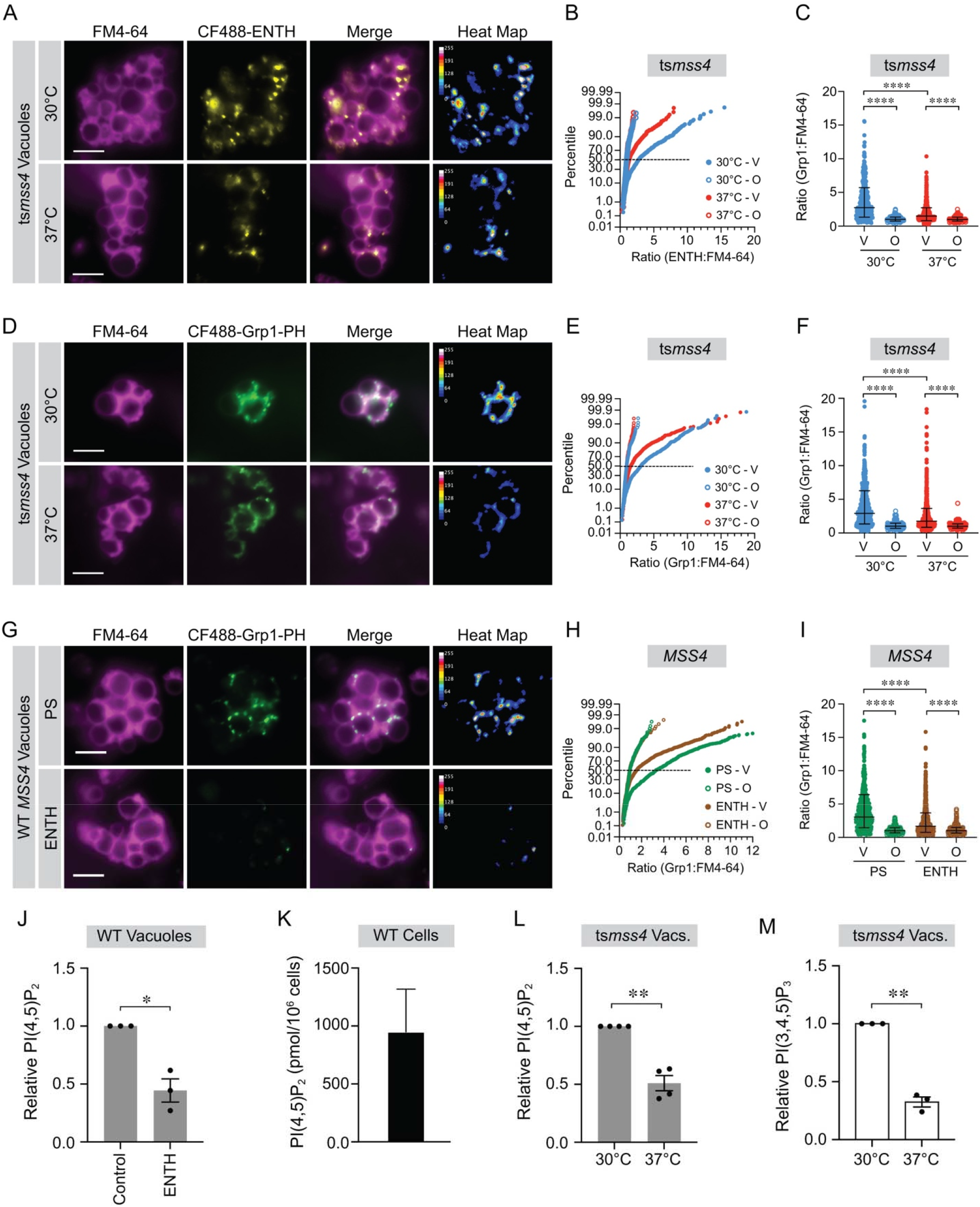
PI(4,5)P_2_ is required for PI(3,4,5)P_3_ production on vacuoles. **(A)** Docking reactions using purified ts*mss4* vacuoles were incubated for 30 min at 30°C or 37°C. PI(4,5)P_2_ was labeled with GST-ENTH and CF488 anti-GST IgG as described in Fig.3. After incubation the reactions placed on ice and labeled with FM4-64 then centrifuged to remove excess antibody and resuspended in PS buffer. Reactions were mounted on glass slides as described and observed by fluorescence microcopy. Scale bars: 5 µm. ENTH/PI(4,5)P_2_ labeling was pseudocolored yellow to distinguish from PI(3,4,5)P_3_ labeling. Fluorescence channels are shown separately as well as overlayed (Merge). Heat maps of ENTH labeling were generated to show relative intensities with a calibration scale. **(B)** Cumulative distribution plot of pooled experiments (n=3) (30°C-V = 519; 30°C-O = 199; 37°C-V = 478; 37°C-O = 217). **(C)** Pooled data shown as a scatter plot with geometric means and their geometric standard deviations. Significance was determined using one-way ANOVA for multiple comparisons and Tukey’s test for individual *p*-values. \*\*\*\**p*<0.0001. **(D)** Docking reactions using purified ts*mss4* vacuoles were incubated for 30 min at 30°C or 37°C and labeled with GST-Grp1-PH and CF488 anti-GST IgG as described. Reactions were prepared for fluorescence microscopy as described above. Scale bars: 5 µm. **(E)** Cumulative distribution plot of pooled experiments (n=3) (30°C-V = 721; 30°C-O = 240; 37°C-V = 667; 37°C-O = 231). **(F)** Scatter plot of data from panel G with geometric means and geometric standard deviations. Significance was determined using one-way ANOVA for multiple comparisons and Tukey’s test for individual *p*-values. \*\*\*\**p*<0.0001. **(G)** Docking reactions with WT (*MSS4*) vacuoles incubated with ENTH lacking GST to physically block PI(4,5)P_2_ and labeled with GST-Grp1-PH to track PI(3,4,5)P_3_. Images. **(H)** Cumulative distribution plots as above (Control-V = 656; Control-O = 294; ENTH-V = 894; ENTH-O = 398). **(I)** Scatter plots of pooled data shown in panel I. Error bars represent geometric means ± geometric standard deviations. Significance was determined using one-way ANOVA for multiple comparisons and Tukey’s test for individual *p*-values. \*\*\*\**p*<0.0001. **(J)** ELISA quantitation of WT vacuole PI(4,5)P_2_ in the presence or absence of ENTH. **(K)** ELISA quantitation of WT cellular PI(4,5)P_2_. **(L)** ELISA quantitation of ts*mss4* vacuole PI(4,5)P_2_ at 30°C and 37°C with or without ENTH. **(M)** ELISA quantitation of ts*mss4* vacuole PI(3,4,5)P_2_ at 30°C and 37°C.

To see the effect on lowering PI(4,5)P_2_ levels on PI(3,4,5)P_3_ labeling, we used ts*mss4* vacuoles as described above and labeled membranes with CF488-Grp1-PH to detect PI(3,4,5)P_3_. When incubated at 30°C, we observed strong CF488-Grp1-PH labeling; however, when incubated at 37°C, there was a sharp reduction in CF488-Grp1-PH fluorescence, indicating that reducing PI(4,5)P_2_ availability had a negative effect on the production of PI(3,4,5)P_3_ **(Fig. 7D-F).** To further examine if PI(4,5)P_2_ availability inhibited PI(3,4,5)P_3_ production, we used 600 nM ENTH (without the GST tag) to sequester PI(4,5)P_2_ from Vps34 kinase activity. This approach reduced PI(3,4,5)P_3_ labeling in a manner comparable to experiments using ts*mss4* inactivation **(Fig. 7G-I)**. Together, these data signal that PI(4,5)P_2_ was a precursor of PI(3,4,5)P_3_ production in *S. cerevisiae*.

### ts*mss4* inactivation and PI(3,4,5)P_3_ quantitation

Although microscopy experiments showed that PI(4,5)P_2_ availability was required for PI(3,4,5)P_3_ production, a quantitative measurement would strengthen our conclusions. For this we used quantitative ELISAs for detecting each lipid. First, we measured PI(4,5)P_2_ levels from WT Mss4 containing vacuoles. This was performed in the presence or absence of ENTH to compete for detection. We found that blocking detection with ENTH reduced it by ∼50% **(Fig. 7J)**. The data was normalized to 1 to account for day-to-day variability in extraction efficiency. To compare vacuolar PI(4,5)P_2_ to the total amount in cells, we extracted lipids from whole WT cells as described. We found that PI(4,5)P_2_ levels ranged from 440 to 1,650 pmol/10^6^ cells with an average of 1,240 pmol/10^6^ cells **(Fig. 7K)**. This difference exceeded what was seen with PI(3,4,5)P_3_ distribution between vacuoles and whole cells. Finally, we extracted lipids from isolated ts*mss4* vacuoles incubated at 30°C or 37°C to quantitate PI(4,5)P_2_ production. This showed that incubating ts*mss4* vacuoles at 37°C reduced PI(4,5)P_2_ concentrations by ∼40% (32 to 19 pmol/10^6^ cells) **(Fig. 7L)**. Ultimately, we wanted to see how inactivating PI(4,5)P_2_ production affected PI(3,4,5)P_3_ levels. We found that inactivating ts*mss4* at 37°C reduced PI(3,4,5)P_3_ levels by 60% compared to vacuoles incubated at 30°C. **(Fig. 7M)**. It is important to remember that ts*mss4* inactivation only affects new lipid production. Thus, preexisting PI(4,5)P_2_ could still be converted to PI(3,4,5)P_3_ Together, these data demonstrate that PI(4,5)P_2_ serves as a substrate for PI(3,4,5)P_3_ production by Vps34.

### PI(3,4,5)P_3_ affects GFP-Ypt7 and HOPS enrichment at vertex domains

Previous studies have shown that regulatory lipids are essential for the formation of vertex microdomains, thus we asked if PI(3,4,5)P_3_ could also affect these platforms (Fratti et al., 2004; Zhang et al., 2024; Sasser et al., 2012). To do this, we used vacuoles containing GFP-Ypt7 and measured its vertex enrichment when PI(3,4,5)P_3_ was sequestered by Grp1-PH. The untreated control showed enriched GFP-Ypt7 at vertex sites (arrows) while outer edge membranes were weakly stained (arrow heads) **(Fig. 8A-C)**. Treating vacuoles with Grp1-PH prevented GFP-Ypt7 from accumulating at vertices, resulting in its dispersion throughout the membrane as seen by the increased staining at outer edges. The difference is clearest in the heat map where outer edges labeled more when Grp1-PH was added. This demonstrated that PI(3,4,5)P_3_ availability was essential for proper mobilization of Ypt7 to the sites of fusion. To confirm that endogenous PI(3,4,5)P_3_ availability affected GFP-Ypt7 distribution, we used 250 µM C8-PI(3,4,5)P_3_ as a competitive inhibitor. Similar to the effects of Grp1-PH, we found that adding C8-PI(3,4,5)P_3_ led to the redistribution of GFP-Ypt7 throughout membranes **(Fig. 8D-F)**. These results demonstrated that PI(3,4,5)P_3_ was needed for the assembly of vertex microdomains.

**Figure 8.**
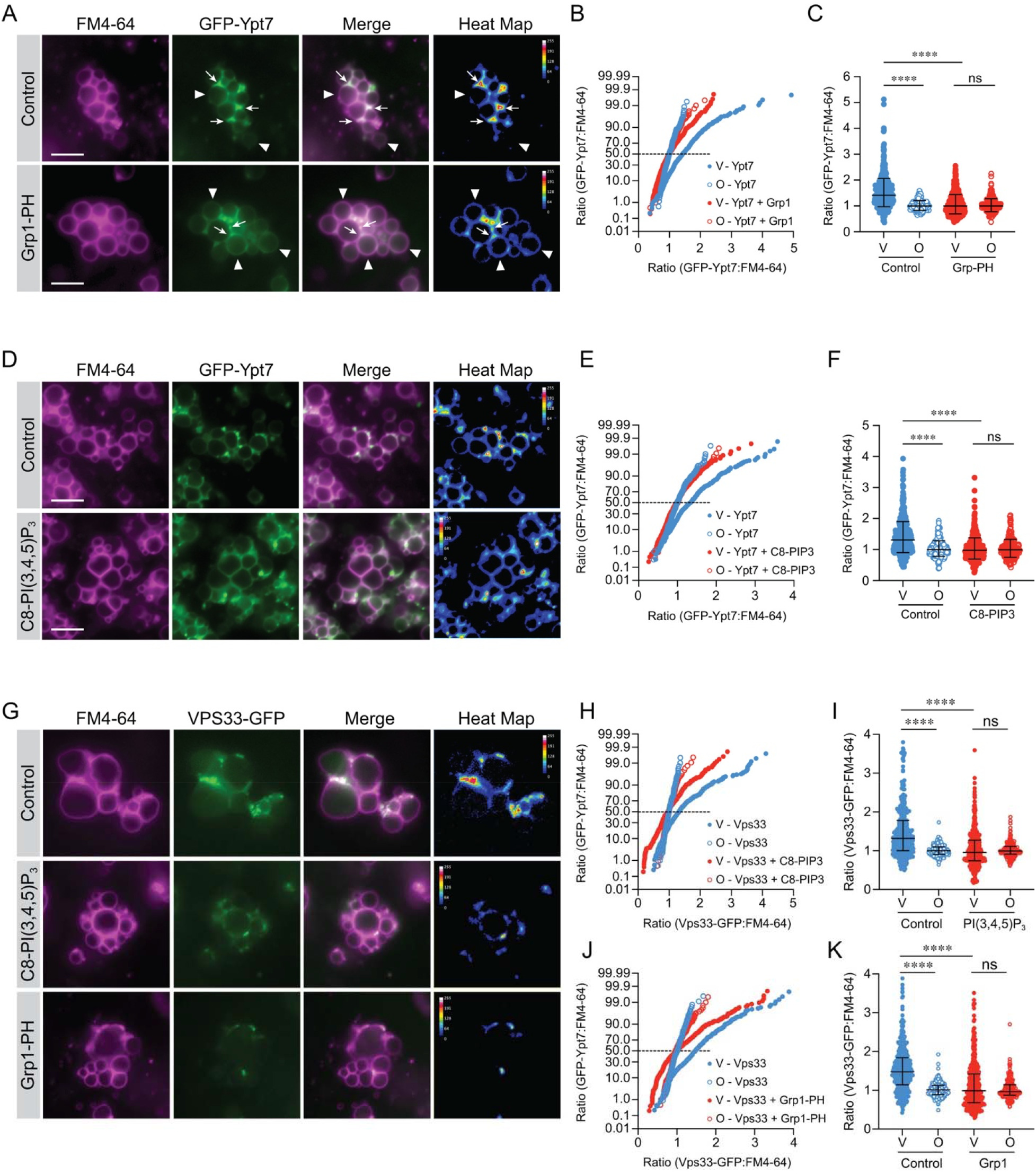
Grp1-PH blocks vertex enrichment of GFP-Ypt7 and Vps33-GFP. **(A)** Docking reactions with vacuoles containing GFP-Ypt7 were treated with unlabeled 1.5 µM Grp1-PH to inhibit fusion or buffer and incubated for 30 min at 27°C. At the end of the incubation period, reactions were prepared for fluorescence microscopy. **(B)** Cumulative distribution plot of pooled ratiometric fluorescence intensities of vertices (V) and outer edge (O) in panel A (n=3) (Control C-V = 438; Control C-O = 171; Grp1-PH C-V = 498; Grp1-PH C-O = 204). **(C)** Pooled data shown as a scatter plot with geometric means and their geometric standard deviations. **(D)** Docking reactions with GFP-Ypt7 vacuoles were treated with 250 µM C8-PI(3,4,5)P_3_ to inhibit fusion or buffer and incubated for 30 min at 27°C. **(E)** Cumulative distribution plot of pooled ratiometric fluorescence intensities of vertices (V) and outer edge (O) in panel D (n=3) (Control C-V = 593; Control C-O = 308; C8-PI(3,4,5)P_3_ C-V = 423; C8-PI(3,4,5)P_3_ C-O = 220). **(F)** Quantitation of ratiometric fluorescence intensities of vertices (V) and outer edge (O) in panel E. Data points were pooled from multiple experiments as described. Error bars represent geometric means ± geometric SD (n=3). **(G)** Docking reactions were performed with vacuoles with Vps33-GFP in the presence buffer (control), 250 µM C8-PI(3,4,5)P_3_ or 1.5 µM Grp1-PH At the end of the incubation period, reactions were processed for fluorescence microscopy as described above. **(H)** Cumulative distribution plot of pooled ratiometric fluorescence intensities of control and C8-PI(3,4,5)P_3_-treated vacuoles. Points represent each vertex (V) and outer edge (O) measurement for the experiments (n=3) (Control C-V = 369; Control C-O = 203; C8-PI(3,4,5)P_3_ C-V = 489; C8-PI(3,4,5)P_3_ C-O = 216). **(I)** Quantitation of ratiometric fluorescence intensities of vertices (V) and outer edge (O) in panel H. Data points were pooled from multiple experiments as described. Error bars represent geometric means ± geometric SD (n=3). **(J)** Cumulative distribution plot of pooled experiments of control and Grp1-PH treated vacuoles (n=3) (Control C-V = 429; Control C-O = 226; Grp1-PH C-V = 477; Grp1-PH C-O = 202). **(K)** Quantitation of ratiometric fluorescence intensities of vertices (V) and outer edge (O) in panel J. Error bars represent geometric means ± geometric SD. Significance was determined using one-way ANOVA for multiple comparisons and Tukey’s post-hoc test for individual *p*-values. \*\*\*\**p*<0.0001; ns – not significant. Scale bars: 5 µm.

To see if other vertex components were affected by PI(3,4,5)P_3_, we used vacuoles that contained Vps33-GFP, a HOPS subunit. Vps33-GFP localization to vertices has been shown to be sensitive to the lipid binding probes that target PI3P, PI(4,5)P_2_, DAG and ergosterol (Fratti et al., 2004). Moreover, Vps33 can be released from vacuoles when PI3P, PI4P and PI(4,5)P_2_ are sequestered (Stroupe et al., 2006). Here, we tested the effects of C8-PI(3,4,5)P_3_ and Grp1-PH on Vps33-GFP distribution. Untreated vacuoles contained enriched Vps33 at vertices as expected **(Fig. 8G-I)**. However, Vps33-GFP vertex enrichment was sharply reduced by C8-PI(3,4,5)P_3_ **(Fig. 8G-I)** and Grp1-PH **(Fig. 8G, J-K)**, further demonstrating that free PI(3,4,5)P_3_ was required for normal vertex assembly.

### PI(3,4,5)P_3_ affects trans-SNARE pairing

While blocking or altering PI(3,4,5)P_3_ affected the Ypt7-dependent tethering stage, the reduction did not account for the overall inhibition of fusion. Thus, it was likely that vertices that formed could be blocked from fusing after tethering. This was consistent with the gain of resistance experiments for targeting PI(3,4,5)P_3_, which were later compared to the tethering/GDI curve. The first known event downstream of tethering is the formation of trans-SNARE complexes between partner membranes. To examine trans-SNARE pairing, we used two types of vacuoles **(Fig. 9A)** (Collins and Wickner, 2007; Jun and Wickner, 2007). One type lacked the R-SNARE Nyv1 and expressed the Qa-SNARE syntaxin ortholog Vam3 with an internal calmodulin binding peptide between the N-terminal helical H_abc_ domain and the SNARE domain (CBP-*VAM3 nyv1*Δ). The second type contained Nyv1 and unmodified Vam3 (*VAM3 NYV1*). The formation of trans-SNARE complexes was detected when Nyv1 co-isolated with CBP-Vam3. We found that under control conditions CBP-Vam3 indeed paired with Nyv1 from partner membranes, as well as the HOPS subunits Vps33 and Vps18 **(Fig. 9B-C)**. As a negative control, we used NEM to inhibit SNARE priming and thus prevented downstream trans-SNARE interactions. As expected, NEM treatment abolished Nyv1 co-isolation with CBP-Vam3 but had no effect on Vam3-HOPS interactions **(Fig. 9B and C, lanes 7 vs 8)**.

**Figure 9.**
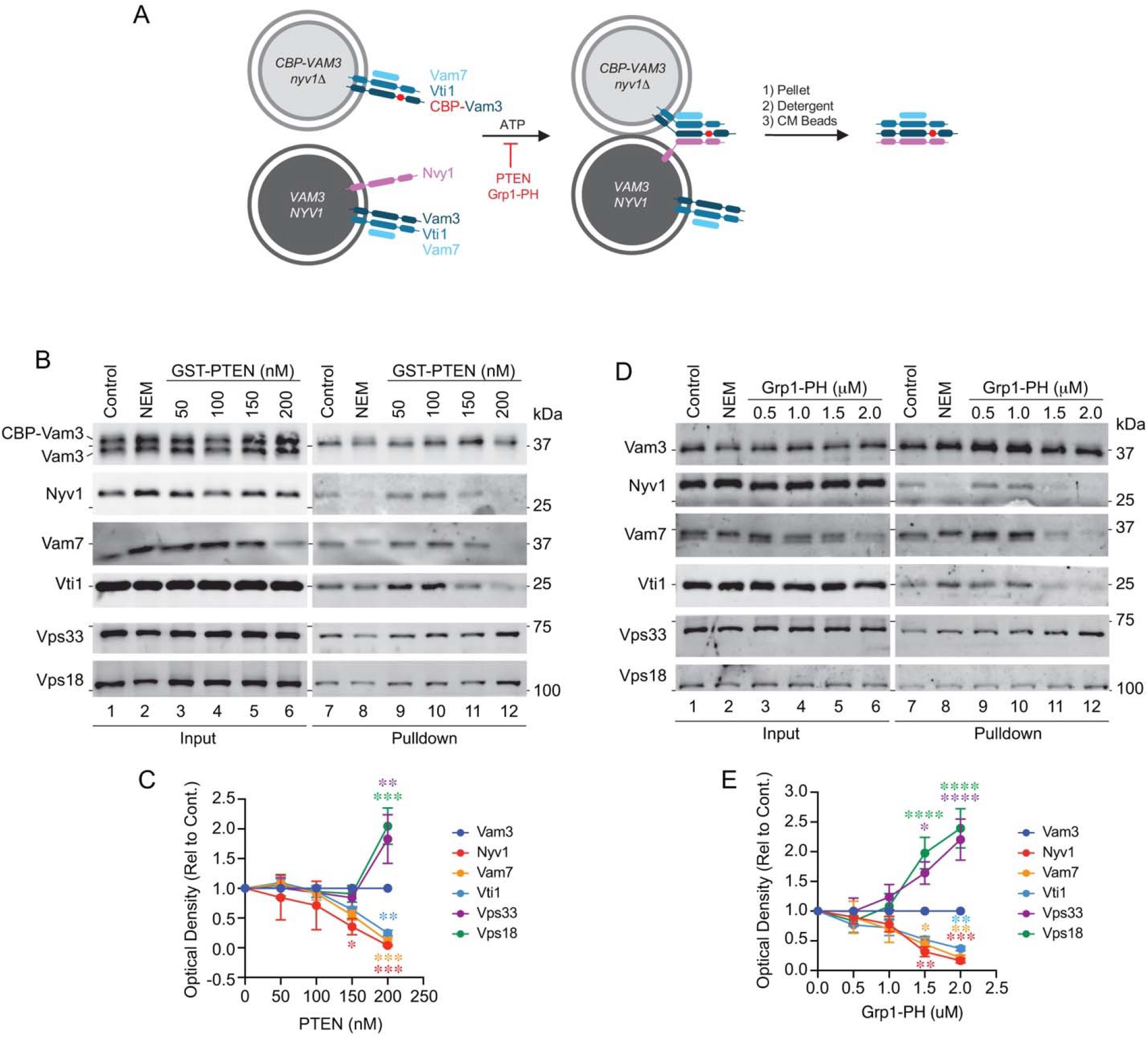
PI(3,4,5)P3 affects trans-SNARE pairing. **(A)** Trans-SNARE isolation schematic. **(B)** Fusion reactions were treated with buffer (Control), 2 mM NEM, or GST-PTEN at the indicated concentrations. Reactions were incubated for 60 min at 27°C. Protein complexes containing CBP-Vam3 were isolated with calmodulin beads. Proteins were resolved by SDS-PAGE and processed for immunoblotting for Vam3, Nyv1, Vam7, Vti1, Vps33 and Vps18. **(C)** Quantitation of proteins bound to CBP-Vam3 in the presence of PTEN. Data represents mean and SEM (n=3). Significance was determined using one-way ANOVA for multiple comparisons and Šidák’s test for pairwise comparisons between CBP-Vam3 and specified proteins for individual *p*-values. **(D)** Reactions were treated with buffer (Control), 2 mM NEM, or GST-Grp1-PH at the indicated concentrations and processed as described above. **(E)** Quantitation of CBP-Vam3 interactions as described above. Data represents mean and SEM (n=3). \**p*<0.05, \*\**p*<0.01, \*\*\**p*<0.001, \*\*\*\**p*<0.0001.

To test the role of PI(3,4,5)P_3_ in trans-SNARE pairing, we used a concentration curve of PTEN and found that it blocked CBP-Vam3 pairing with Nyv1 starting at 150 nM with a further decrease at 200 nM. The inhibition of trans-SNARE complex formation could be due to two factors. First, the Qb-SNARE Vti1 was lost from the CBP-Vam3 complexes when PTEN was at ≥ 150 nM, even though it was present in the input **(Fig. 9B, lane 6 vs 12, and C)**. This was significant because the Q-SNAREs form stable complexes prior to binding their respective R-SNARE in trans (Jun et al., 2006; Pribicevic et al., 2024; Fasshauer and Margittai, 2004) and suggested that PI(3,4,5)P_3_ affected SNARE pairing at more than one point. Second, Vam7 was depleted in the input membrane fraction and completely absent from CBP-Vam3 complexes when vacuoles were treated with 200 nM PTEN. It is important to note that the first step in trans-SNARE isolation is pelleting membranes, which separates vacuole-bound and unbound proteins. Thus, a release of Vam7 from the vacuole would result in its absence from the input blot.

In addition to SNARE complex formation, CBP-Vam3 also pulled down the HOPS complex under untreated or NEM treated conditions. Surprisingly, when vacuoles were treated with PTEN, we found that it did not inhibit HOPS co-isolation with CBP-Vam3. On the contrary, we discovered that depleting PI(3,4,5)P_3_ enhanced HOPS binding to CBP-Vam3 **(Fig. 9B-C)**. Since tethering precedes docking, it is likely that this represented a pre-trans-SNARE Vam3-HOPS complex, and that PI(3,4,5)P_3_ leads to HOPS-Vam3 separation to allow other SNAREs to interact with Vam3. This is of note because the H_abc_ N-terminal domain of Vam3 alone interacts with HOPS, after which HOPS shifts to bind the SNARE motif bundle of the Q-SNARE complex prior to engaging Nyv1 in trans (Lürick et al., 2015; Laage and Ungermann, 2001).

To confirm the effects of PTEN on trans-SNARE pairing, we repeated the experiment using Grp1-PH. Grp1-PH showed a dose-dependent loss of Vam7 from the input lanes **(Fig. 9D, lanes 3-6)**, which was reflected in pulldown efficiency **(Fig. 9D, lanes 9-12, and E)**. Similarly, there was a gradual loss in Nyv1 and Vti1 in the pulldown lanes without a loss in the input. In addition to its effects on SNARE pairing, Grp1-PH also increased HOPS-Vam3 interactions as seen with PTEN **(Fig. 9D-E).**

These data suggest that the roles of PI(3,4,5)P_3_ include the stabilization of the 3Q-SNARE complex to promote optimal interactions with Nyv1. While not published, we must note that Vam3, Vti1 and Nyv1 have polybasic regions (PBR) adjacent to their transmembrane domains (TMDs) that could interact with acidic lipids (SGD - Saccharomyces Genome Database). A similar juxtamembrane PBR is present in Syntaxin1A that interacts with PI(3,4,5)P_3_ to promote clustering and neurotransmitter release (Khuong et al., 2013). Thus, it is possible that vacuolar PI(3,4,5)P_3_ could interact with yeast vacuole SNARE PBRs to promote complex formation and fusion.

### PI(3,4,5)P_3_ binding competition releases Vam7 from membranes

The trans-SNARE pairing experiments showed that Vam7 could be released from membranes when vacuoles were treated with PTEN or Grp1-PH, suggesting that PI(3,4,5)P_3_ was directly or indirectly involved in maintaining Vam7 bound to vacuoles. To further test this, we tracked the bound and unbound fractions of Vam7 when PI(3,4,5)P_3_ was sequestered by Grp1-PH. Fusion reactions were treated with buffer (0 µM) or GST-Grp1-PH at increasing concentrations. After incubation, vacuole-bound and unbound proteins were separated by centrifugation, and the two fractions were examined by immunoblotting. As seen previously, soluble/peripheral proteins such as HOPS, actin, and Vam7 were seen in both bound and unbound fractions (Fratti et al., 2004; Stroupe et al., 2006; Lawrence et al., 2014) **(Fig. 10A-B)**. In contrast, the membrane-anchored proteins, Nyv1 and Ypt7, were only seen in the bound fraction. Treatment with GST-Grp1-PH strongly affected Vam7, as its release was the most responsive to GST-Grp1-PH addition **(Fig. 10A lanes 5 and 11)**. This was reminiscent of a previous study showing that Vam7 was released in the presence of the lipid binding domains FYVE, ENTH and C1b, which bind PI3P, PI(4,5)P_2_ and DAG, respectively (Fratti et al., 2004). HOPS subunits were also affected by varying degrees. Vps18 was moderately released at 2 µM GST-Grp1-PH, whereas Vps33 release was more pronounced and occurred at lower GST-Grp1-PH levels. Vps33 release has been shown to be strongly affected by FYVE and ENTH, as well as the PI4P binding domain Fapp1-PH (Stroupe et al., 2006). We also tested C8-PI(3,4,5)P_3_ at 250 µM which inhibits fusion. Unlike Grp1-PH, C8-PI(3,4,5)P_3_ had no effect on Vam7 or HOPS binding. This could be due to differences in binding affinities where C8-PI(3,4,5)P_3_ cannot compete with full length lipid. It is also possible that C8-PI(3,4,5)P_3_ affected fusion by altering vertex microdomains without displacing Vam7.

**Figure 10.**
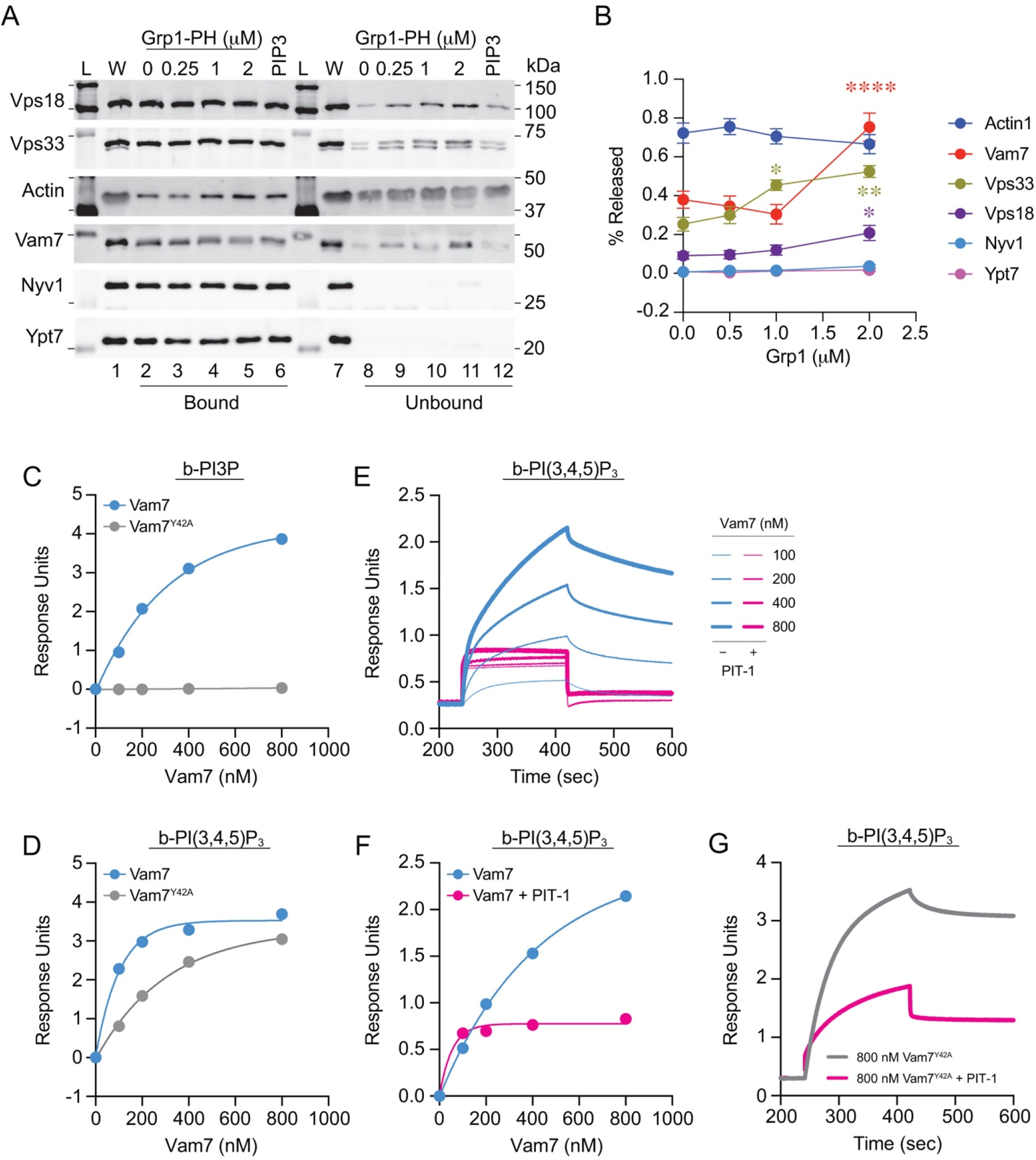
Vam7 Interacts with PI(3,4,5)P_3_. **(A)** Fusion reactions were treated with buffer (Control) or GST-Grp1-PH at the indicated concentrations. Reactions were incubated for 60 min at 27°C. Membrane-bound and unbound proteins were separated by centrifugation (16,000 g, 10 min, 4°C) into unbound and bound fractions. Pellets were resuspended in PS buffer to match the starting volumes. Bound and unbound fractions were mixed with SDS-loading buffer and processed for immunoblotting for Actin, Nyv1, Vam7, Ypt7, Vps33 and Vps18. **(B)** Quantitation of proteins released from vacuoles in the presence of Grp1-PH. Data represents mean and SE (n=4). Significance was determined using one-way ANOVA for multiple comparisons and Šidák’s test for pairwise comparisons for individual *p*-values. \*\*\*\**p*<0.0001. L, ladder; W, whole reaction. **(C-D)** BLI binding curves of b-PI3P (C) or b-PI(3,4,5)P_3_ (D) in response units versus protein concentrations of Vam7 and Vam7^Y42A^. **(E-F)** BLI binding curves of PIT-1 (200 µM) inhibition of b-PI(3,4,5)P_3_ binding Vam7. **(G)** BLI sensorgrams of 800 nM Vam7^Y42A^ to b-PI(3,4,5)P_3_ with and without 200 µM PIT-1.

### Vam7 binds PI(3,4,5)P_3_

The disproportionate release of Vam7 suggested that PI(3,4,5)P_3_ could directly affect the SNARE in addition to regulating vertex microdomain assembly/maintenance. Vam7 binds to vacuole membranes through its PX domain that interacts with PI3P, and mutating Tyr42 to Ala blocks PI3P binding, severely impairing fusion (Boeddinghaus et al., 2002; Fratti and Wickner, 2007; Cheever et al., 2001). The ability of Vam7^Y42A^ to support any fusion was attributed to its interactions with SNAREs and HOPS without the aid of PI3P. That said, it could be that Vam7^Y42A^ interacted with other lipids, including PI(3,4,5)P_3_, either specifically or non-specifically. The Vam7 PX domain has a polybasic region outside of the PI3P binding site that could accommodate interactions with anionic lipids. For instance, we previously showed that Vam7 bound to PA and PI5P by liposome flotation, although binding affinities were not determined (Miner et al., 2016). Separately, we found that PA binding was very weak as measured by surface plasmon resonance, microscale thermophoresis and BLI (Sparks et al., 2019, 2022; Calderin et al., 2025). Thus, we can surmise that Vam7 could either have a second lipid-binding site or that its single site is promiscuous, allowing multiple lipids to bind with variable affinities. The former is in accord with a study showing that many PX domains have a second lipid binding site (Chandra et al., 2019).

Here we compared PI3P and PI(3,4,5)P_3_ binding using streptavidin-coated BLI probes bound to biotinylated lipids. These were incubated with GST-Vam7 and GST-Vam7^Y42A^ at different concentrations to measure binding affinities. In Fig.10 we show curves of response units versus protein concentration. This demonstrated that Vam7 bound to b-PI3P whereas Vam7^Y42A^ failed to bind, as expected **(Fig. 10C)**. In comparison, we found that both Vam7 and Vam7^Y42A^ bound to b-PI(3,4,5)P_3_ **(Fig. 10D)**. The binding of Vam7^Y42A^ to PI(3,4,5)P_3_ was also consistent with the idea of a second binding site. Alternatively, other residues in the same binding pocket might have engaged the D4 and D5 phosphates of PI(3,4,5)P_3_, which were not involved in binding with the D3 phosphate of PI3P. It was also possible that the Y42A mutation weakened PI(3,4,5)P_3_ binding without abolishing it. Future investigation will explore these options.

Finally, we asked if Vam7 binding to b-PI(3,4,5)P_3_ could be blocked. For this we used PIT-1 to inhibit PI(3,4,5)P_3_-protein interactions. PIT-1 (200 µM) and Vam7 were pre-incubated together, then transferred to microtiter plate wells for BLI reactions. We found that PIT-1 abolished Vam7 binding to b-PI(3,4,5)P_3_ **(Fig. 10E-F).** This was in accord with the activity of PIT-1 on PI(3,4,5)P_3_ interactions with lipid binding domains. Importantly, PIT-1 reduced Vam7^Y42A^ binding to b-PI(3,4,5)P_3_ **(Fig. 10G)**. This suggests that both WT and Y42A interact with b-PI(3,4,5)P_3_ in a similar manner. While PIT-1 was discovered by looking for inhibitors of PH domains binding to PI(3,4,5)P_3_, it did not exclude non-canonical pairings. This was shown in part by the ability of PIT-1 to block PTEN phosphatase activity **(Fig. 2M)**.

### Co-folding and molecular docking of Vam7 and PI(3,4,5)P_3_

Given the unexpected interaction between Vam7 and PI(3,4,5)P_3_, we set to model differences in how Vam7 binds to PI3P or PI(3,4,5)P_3_. We generated models of full-length Vam7 using its amino acid sequence available at the SGD to binding to short chain PI(3,4,5)P_3_ or PI3P using the Boltz-2 co-folding algorithm, which allows structural and binding affinity predictions benchmarked to approach the accuracy of free-energy perturbation (FEP) calculations at correlating with biophysical measurements (Passaro et al., 2025). This approach allows for all residues of Vam7 to be examined in terms of their effect on binding, which is important because Vam7 autoregulates its binding to PI3P (Miner et al., 2016). In **Fig.11** we show that PI3P and PI(3,4,5)P_3_ are predicted to bind the same pocket in the Vam7 PX domain in co-folding models. Binding to PI3P involved Arg41, Tyr42 and Ser43 interactions with the D3 phosphate with a predicted *K*_D_ of 166 nM **(Fig. 11A and C)**. When binding PI(3,4,5)P_3_, Tyr42 did not interact with the D3 phosphate, whereas Arg41 and Ser43 interacted with both D3 and D4 phosphates. The D5 phosphate interacted with Arg88. The predicted *K*_D_ for PI(3,4,5)P_3_ to full-length Vam7 was 49 nM **(Fig. 11B and C).** The increased predicted affinity for PI(3,4,5)P_3_ is consistent with the BLI experiments described in **Fig. 11**. The absence of Tyr42 interactions with the D3 phosphate is consistent with the ability of Vam7^Y42A^ to bind PI(3,4,5)P_3_.

**Figure 11.**
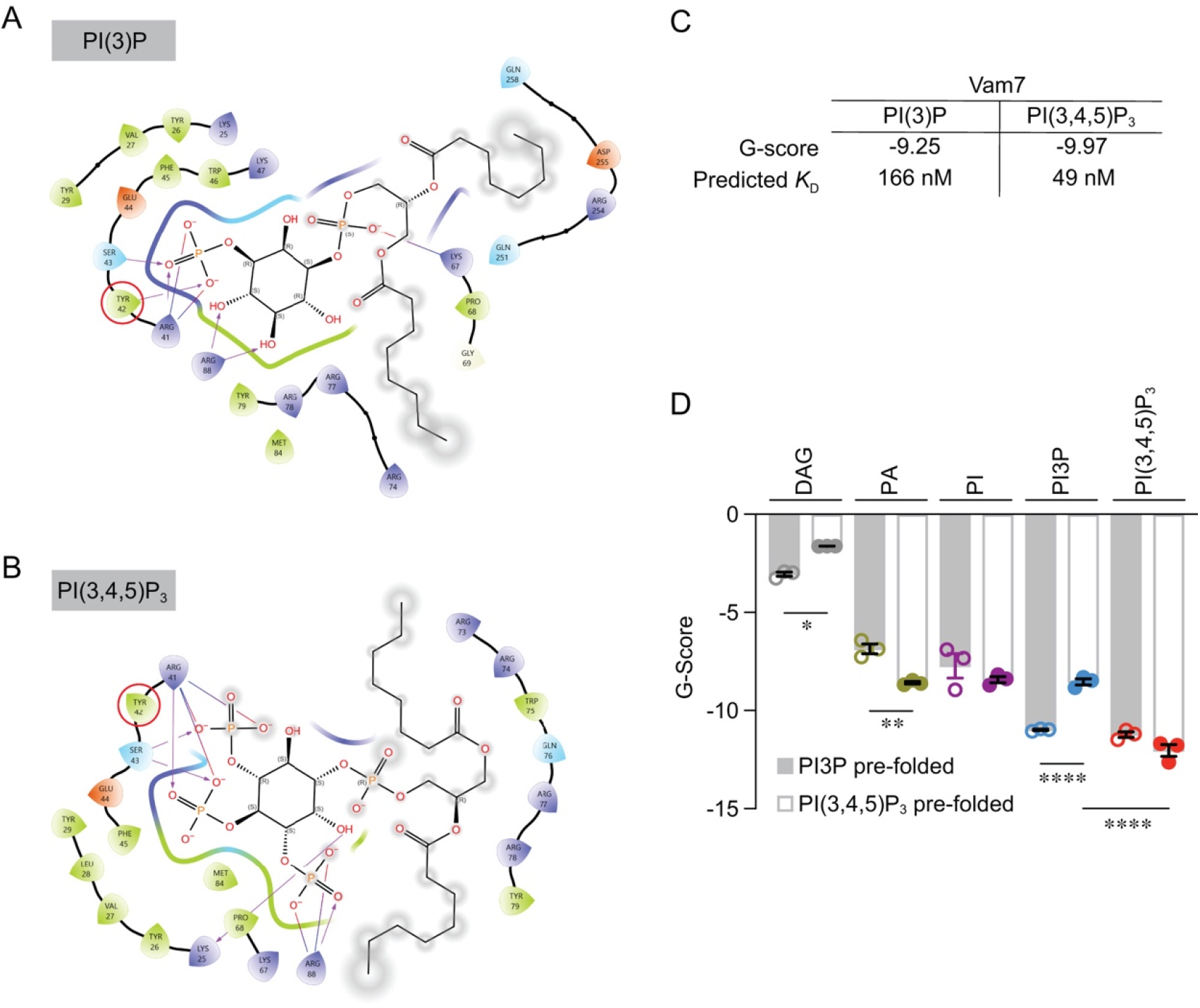
Computational docking of Vam7 and PIP3. **(A)** Ligand interaction diagram of PI3P binding to the PX domain of Vam7 model corresponding to Boltz-2 co-folding predictions for full-length Vam7. Interactions are indicated with arrows showing H-bonding and salt-bridging between Tyr42, Ser43 and Arg41 with the D3 phosphate of PI3P and Arg88 D4 and D5 hydroxyl groups of the inositol ring. **(B)** Ligand interaction diagram of PI(3,4,5)P_3_ binding to the PX domain as in panel A. Interactions are represented with arrows showing H-bonding and salt-bridging between Arg41 and Ser43 with the D3 and D4 phosphates, Arg88 and the D5 phosphate, and Lys25 and the D6 hydroxyl of the inositol ring. **(C)** The predicted g-scores for Glide SP as well as the predicted affinity from Boltz-2 complex predictions between Vam7 and either PI3P or PI(3,4,5)P_3_. **(D)** G-scores from Glide XP for lipid binding to grids centered on Vam7 co-folded with either PI3P or PI(3,4,5)P_3_. Bars represent SEM (n=3). Significance was determined with one-way ANOVA and Tukey’s test for individual p-values. * *p*<0.05, ** *p*<0.01, **** *p*<0.0001.

To further understand binding of short chain PI(3,4,5)P_3_ or PI3P we generated grids around either model, minimized these structures using OPLS4 force field, and introduced a panel of lipids to score binding with Glide XP (Extra-precision) docking (g-score) (Singh et al., 2021). For Vam7 co-folded PI3P model we found that DAG interacted poorly (g-score = -2.98 kcal/mol) whereas PA and PI bound with intermediate values of g-score = -6.8 and -7.2 kcal/mol, respectively (**Fig. 11D, gray bars**). When PI3P and PI(3,4,5)P_3_ were introduced, we found that both had high binding energies of g-score ∼-11 kcal/mol. Interestingly, when Vam7 was co-folded with PI(3,4,5)P_3_ (white bars), the g-score for PI3P was significantly reduced from -11 to -8.4 kcal/mol, whereas binding for PI(3,4,5)P_3_ was not affected.

### WT Vam7 PX domain binds most frequently to PI3P membranes

To determine how the membrane recognition of the Vam7 PX-domain alone was affected by PI3P or PI(3,4,5)P_3_, or by mutations, we performed HMMM simulations of WT and Y42A Vam7 PX on PI3P and PI(3,4,5)P_3_-containing membranes. For this we used the solution NMR structure (https://doi.org/10.2210/pdb1KMD/pdb) of the PX domain (Lu et al., 2002). The simulations contained four independent 250-ns replicas for each of the four protein–lipid configurations, resulting in a total of 16 trajectories and 4.0 µs of aggregate sampling across all conditions. Based on Cα RMSD analysis **(Fig. S2)**, the PX domain showed significant conformational changes in all membrane simulations. Comparing to the smaller RMSD values obtained in solution for the protein (2-2.5 Å; data not shown), the larger changes observed here may reflect membrane-induced conformational rearrangements.

Membrane depth analysis of individual protein residues revealed a clear difference in the membrane pose across the 4 conditions **(Fig. 12A, Table S1)**. WT Vam7 on PI3P (WT+PI3P) membranes formed stable membrane-bound forms in all 4 replicas. In contrast, WT+PI(3,4,5)P_3_, Y42A+PI3P, and Y42A+PI(3,4,5)P_3_ each produced a stably bound form in only one of 4 replicas. Thus, WT+PI3P was the only condition in which the PX domain reproducibly reached the membrane phosphate plane and remained bound for segments of at least 30 ns in all replicas. The reduced occurrence of membrane binding in Y42A+PI3P, compared to WT+PI3P, supports a role for Tyr42 in recognizing PI3P membranes. The per-residue membrane-distance profiles calculated from stably membrane-bound WT+PI3P frames further support this interpretation **(Fig. 12B)**. These profiles show residue-level membrane distances of the 4 stably bound WT+PI3P replicas and provide a structural view of the membrane-bound state under this condition.

**Figure 12:**
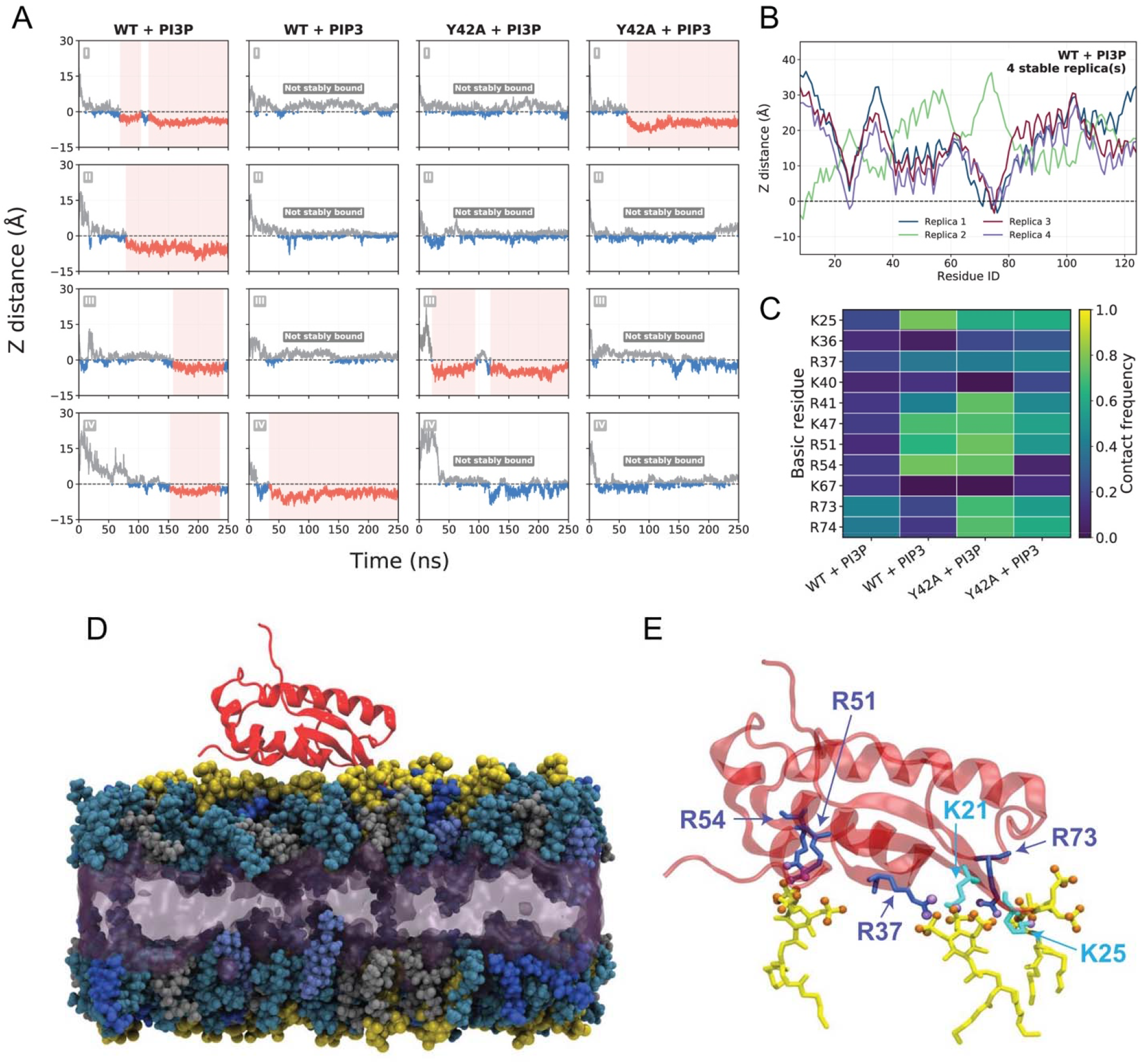
Membrane binding of Vam7 PX domain. **(A)** Membrane-depth analysis of all 16 production MD trajectories. Stably bound intervals are defined as continuous membrane-bound segments lasting at least 30 ns. WT+PI3P achieves stably bound states in all 4 replicas, whereas WT+PIP3, Y42A+PI3P, and Y42A+PIP3 show such state in only one replica each. **(B)** Per-residue membrane-distance profiles calculated from stably bound WT– PI3P frames. The 4 WT+PI3P replicas provide a residue-level view of the membrane-bound configuration associated with the canonical PI3P-recognition state. **(C)** Residue-level Lys/Arg–PIP oxygen contact frequency across all production frames using a 4.0Å cutoff. PIP3-containing systems show broad basic-residue engagement, and Y42A+PI3P retains several basic contacts, indicating that Lys/Arg–PIP contacts alone are not sufficient for the formation of the canonical membrane-bound state. **(D)** Representative protein–membrane system for a PIP3-bound configuration. The membrane contains POPC, POPE, POPS, ergosterol, and PIP3. **(E)** Closed up view of the PIP3-interacting electrostatic interface. Arginines are shown in dark blue, lysines in light blue, and PIP3 lipids in yellow. Basic-residue nitrogen atoms are highlighted in purple, and PIP3 phosphate/ring oxygen atoms in orange, highlighting electrostatic interactions between them.

### PI(3,4,5)P_3_-containing membranes engage more basic residues

Although WT+PI3P was the most frequently membrane binding condition, the Lys/Arg– phosphoinositide oxygen contact heatmap shows that PI(3,4,5)P_3_ membranes engage a broader set of basic residues than PI3P ones **(Fig. 12C)**. These interactions involved a polybasic membrane-binding surface containing K21, K25, R37, R51, R54, and R73 **(Fig. 12C and E, Table S2)**. Y42A+PI3P also retained several Lys/Arg+PI3P contacts, indicating that the mutant could still engage with the negatively charged lipid headgroups. However, these contacts did not produce the same reproducible WT+PI3P geometry. Thus, electrostatic contacts alone were not sufficient for the formation of the canonical PI3P-bound state. While this PBR played a role in lipid interactions, it is important to note that R41 of the canonical PI3P binding pocket had higher contact frequency, in WT PX interactions with PI(3,4,5)P_3_ versus those with PI3P. Higher contact frequencies of R41 were also seen with the interactions of Y42A PX with either lipid.

Analysis of the stably bound ensembles supported the same interpretation **(Supplementary Fig 3)**. UMAP projection of per-residue membrane-distance profiles shows that WT+PI3P contributed the most to the stably bound population. The residue-level distance profiles also showed that three of the WT+PI3P replicas sampled closely related membrane-binding configurations. These data support a model in which WT+PI3P forms a reproducible canonical binding geometry, while PI(3,4,5)P_3_-containing systems are better described by broader, but non-specific electrostatic engagement rather than by one highly recurrent insertion geometry **(Supplementary Fig 4B-C)**.

Taken together, the simulations supported a two-component model. PI3P favors a more reproducible, Tyr42-sensitive canonical binding geometry that is most evident for WT Vam7. PI(3,4,5)P_3_, by contrast, promotes broader basic-residue engagement through its more highly phosphorylated headgroup, allowing both WT and Y42A Vam7 to form electrostatic membrane contacts even when the canonical PI3P-bound geometry was not reproduced.

## Discussion

Our previous studies have examined the roles of phosphoinositides and other signaling lipids on the yeast vacuole fusion cycle (Starr and Fratti, 2019; Miner et al., 2020; Zhang et al., 2022, 2024, 2026). Here, we asked if PI(3,4,5)P_3_ could also affect vacuole fusion. PI(3,4,5)P_3_ was thought not to exist in *S. cerevisiae* due to the absence of orthologs of the class I PI3K p110 or its regulator p85. However, others found that the fission yeast *S. pombe* could make PI(3,4,5)P_3_ using the class III PI3K Vps34 (Mitra et al., 2004). Vps34 – also known as PIKC3C/hVps34 – makes PI3P throughout eukarya, yet there is only one example of it making PI(3,4,5)P_3_. In *S. cerevisiae*, Vps34 is responsible for many pathways including macroautophagy, endolysosomal trafficking and vacuole homotypic fusion (Reidick et al., 2017). Based on this, we asked if *S. cerevisiae* Vps34 could make PI(3,4,5)P_3_ on vacuoles. In the end, we showed that Vps34 can indeed make PI(3,4,5)P_3_ from PI(4,5)P_2_ and that it serves multiple functions in the fusion cycle.

In this study, we showed that PI(3,4,5)P_3_ was involved in multiple stages of the fusion pathway, as it affected the formation of vertex membrane microdomains, retaining Vam7 bound to vacuoles and the formation of trans-SNARE complexes. Initially, we used C8-PI(3,4,5)P_3_ as a competitive inhibitor for any unidentified interactions with a potential endogenous long chained lipid counterpart. In comparison with other C8 lipids, C8-PI(3,4,5)P_3_ was the most potent in blocking fusion, suggesting that an endogenous source may exist and that it interacts with components of the fusion machinery. Expanding on this notion, we examined how blocking endogenous PI(3,4,5)P_3_ with the Grp1-PH binding domain or modifying it with the phosphatase PTEN would compare with the initial finding. We found that both Grp1-PH and PTEN blocked fusion with the same timing pattern as C8-PI(3,4,5)P_3_.

While these findings were compelling, it was possible that there was crosstalk with other lipids, most importantly PI3P. As part of testing for this, we examined how well Grp1-PH discriminated for PI(3,4,5)P_3_. BLI experiments showed that Grp1-PH strongly bound PI(3,4,5)P_3_, while it failed to interact with PI3P. Importantly, the Grp1-PH^K273A^ point mutant with reduced affinity for PI(3,4,5)P_3_ failed to bind the lipid and failed to inhibit fusion. Another important distinction between targeting PI(3,4,5)P_3_ and PI3P is the gain of resistance timing. Blocking PI3P with the FYVE domain or the PX domain from Vam7 inhibits fusion with the same half-time curve as GDI, RDI (Rho GDI), and antibodies against vacuolar SNAREs, HOPS, and Ypt7 in what is referred to as the docking curve (Boeddinghaus et al., 2002; Haas et al., 1995; Eitzen et al., 2000; Ungermann et al., 1998a; Eitzen et al., 2001; Ungermann et al., 1999; Mayer and Wickner, 1997). In comparison, Grp1-PH, PTEN, and C8-PI(3,4,5)P_3_ blocked after the docking curve, similar to the inhibition curves of the actin disruptor Latrunculin, the PI(4,5)P_2_ binding factors ENTH and MARCKS effector domain, and C8-PI(3,5)P_2_ (Eitzen et al., 2002; Collins et al., 2005; Miner et al., 2019). This experiment cannot further separate effects as targeting PI(3,4,5)P_3_ blocks trans-SNARE pairing, whereas targeting PI(3,5)P_2_ blocks after SNARE complex formation and before hemifusion (Miner et al., 2019).

### PI(3,4,5)P_3_ enrichment

We have previously visualized lipids on vacuoles using epifluorescence microscopy and labeled protein domains (Fratti et al., 2004; Karunakaran et al., 2012; Sasser et al., 2012, 2013; Miner et al., 2019; Zhang et al., 2024). Typically, lipid binding protein domains are recombinantly expressed and conjugated to fluorophores, which allows the localization of lipids including PI3P, PI(4,5)P_2_, PI(3,5)P_2_, and DAG (Fratti et al., 2004; Miner et al., 2019). Here, we used fluorescent anti-GST antibody to track GST-Grp1-PH. By using the compound reporter, we were able to clearly label PI(3,4,5)P_3_ on vacuoles and determined that it was enriched at the vertices of docked membranes. PI(3,4,5)P_3_ labeling was blocked by competion with C8-PI(3,4,5)P_3_ but not C8-PI(4,5)P_2_. Importantly, Grp1-PH labeling was also blocked when endogenous PI(3,4,5)P_3_ was eliminated with PTEN. The role of PTEN and vacuolar PI(3,4,5)P_3_ was complemented by using vacuoles from a strain lacking the PTEN ortholog Tep1 (Heymont et al., 2000). This shows that just as the addition of PTEN reduced labeling, eliminating *TEP1* augmented PI(3,4,5)P_3_ enrichment.

### PI(3,4,5)P_3_ production

Using the CF488-Grp1-PH labeling method combined with ratiometric quantitation showed PI(3,4,5)P_3_ accumulation and depletion in vitro with isolated vacuoles. This raised the question of how it was made. As mentioned above, others had showed that *S. pombe* could make PI(3,4,5)P_3_ in a Vps34-dependent manner and that the PTEN ortholog Ptn1 could degrade it (Mitra et al., 2004). To examine the role of Vps34 in *S. cerevisiae* PI(3,4,5)P_3_ production, we first used the specific class III PI3K inhibitor SAR405 (Ronan et al., 2014). This strongly reduced Grp1-PH staining and detectable amounts PI(3,4,5)P_3_ on vacuoles, which was in accord with the role of Vps34 in the production of PI(3,4,5)P_3_. Similar results were found with fixed and spheroplasted cells. To show a direct role for Vps34, we used a temperature sensitive mutant. Vacuoles from ts*vps34* cells were used in docking experiments as typically performed, except that a second set of reactions were incubated at the non-permissive temperature (37°C). Inactivating ts*vps34* reduced PI(3,4,5)P_3_ staining similar to what was seen with SAR405. It was not a complete block as some PI(3,4,5)P_3_ was present at vertices. Alternatively, the detectable lipid could have been there before the experiment was performed. This was also seen when ts*vps34* vacuoles were used to extract lipids and quantify PI(3,4,5)P_3_ by ELISA. Heat inactivation reduced PI(3,4,5)P_3_ by ∼50%, which was consistent with the effect of treating with PTEN.

To further illustrate the role of Vps34 in PI(3,4,5)P_3_ production, we expressed GST-mCherry-Grp1-PH in ts*vps34* cells and looked for vacuole staining. This showed that mCherry-Grp1-PH labeled vacuoles and the plasma membrane when cells were grown at 30°C, whereas labeling was reduced at 37°C. A downside to using live cells is that we cannot synchronize fusion events to track vertex enrichment during docking or visualize trans-SNARE pairing. These cells looked different than the fixed spheroplasted ones, which showed strong plasmalemma labeling and invisible vacuole pools. Fixing with formaldehyde permeabilizes cells allowing leakage of cytoplasmic Grp1-PH, which eliminated the haziness seen in live cells. Another difference could be due to the excess of GST-Grp1 used that could compete for PI(3,4,5)P_3_-bound proteins to generate a stronger signal that would mask vacuole staining. Nevertheless, inactivating Vps34 reduced staining in every condition tested.

ELISA measurements of whole cells and vacuolar PI(3,4,5)P_3_ showed that the plasmalemma has ∼31 times more PI(3,4,5)P_3_ than the vacuole pool. This could be due to a natural difference as seen in mammalian cells, or it could be the result of vacuole isolation itself. The isolation of vacuoles is a slow process that involves cell wall degradation, osmotic lysis and flotation through a density gradient. Thus, it is not unlikely that a significant fraction of lipid was lost or degraded during isolation, leading to a lower readout compared to endogenous levels. That said, the vacuoles remained biologically active and could produce fresh PI(3,4,5)P_3_ under fusogenic conditions.

### PI(4,5)P_2_ is the direct substrate for PI(3,4,5)P_3_

Our data using ts*vps34* showed that PI(3,4,5)P_3_ production on vacuoles was dependent on the class III PI3K; however, it did not show that the direct substrate was PI(4,5)P_2_. To address this, we used the ts*mss4* mutant to block PI(4,5)P_2_ production from PI4P. Heat inactivation of Mss4 reduced PI(4,5)P_2_ levels as measured by ELISA as well as fluorescence microscopy. Importantly, isolated ts*mss4* vacuoles incubated at the non-permissive temperature also showed reduced PI(3,4,5)P_3_. Finally, we showed that using the ENTH domain (lacking GST) to sequester PI(4,5)P_2_ away from Vps34 kinase activity blocked PI(3,4,5)P_3_ labeling of WT vacuoles. Taken together, we conclude that PI(3,4,5)P_3_ is made by Vps34 using PI(4,5)P_2_ as a substrate. It is worth noting that preliminary ELISA experiments with increased ATP, longer incubation times and the addition of 3-nitrocoumarin to inhibit PLC activity resulted in increased vacuolar PI(3,4,5)P_3_ detected. This further supports the notion that PI(4,5)P_2_ levels directly affect PI(3,4,5)P_3_ production.

While more convoluted pathways could be drawn to link Vps34 and PI(3,4,5)P_3_, it is most likely that the more direct pathway of adding a third phosphate to PI(4,5)P_2_ was used. For example, one could consider that Vps34 makes PI3P that is converted by Fab1 to PI(3,5)P_2_, which would require the PI 4-kinase Lsb6 to create PI(3,4,5)P_3_ (Han et al., 2002; Gary et al., 1998). However, this would exclude the need for Mss4 and PI(4,5)P_2_ production. Moreover, ENTH domain would need to compete for Lsb6 activity. We also tested the effect of blocking Fab1 with Apilimod and found that it had no effect on PI(3,4,5)P_3_ vacuole labeling **(Supplemental Fig. 4)**. Importantly, Cheever et al. showed that deleting Fab1 did not affect Vam7 association to vacuoles, indicating that PI3P and PI(3,4,5)P_3_ levels were not affected (Cheever et al., 2001). Another pathway would require Vps34 to add a phosphate to PI4P to make PI(3,4)P_2_, after which Mss4 would add the final phosphate at D5. Again, blocking PI(4,5)P_2_ with ENTH would not compete for producing PI(3,4,5)P_3_.

### PI(3,4,5)P_3_, Vam7, and vertex microdomains

Stage specific experiments showed that both Ypt7-dependent tethering and trans-SNARE pairing required PI(3,4,5)P_3_. During docking and tethering, vacuoles become tightly apposed with flattened discs of membrane forming a “fusion synapse” of sorts. At the edge of the discs where membranes come into contact is the vertex ring composed of separate vertex microdomains that become enriched in ergosterol, DAG, PI3P and PI(4,5)P_2_ along with the proteins that drive fusion including SNAREs, Ypt7, HOPS and actin (Wang et al., 2002, 2003; Eitzen et al., 2002; Karunakaran et al., 2012; Fratti et al., 2004). Here, we documented that interfering with PI(3,4,5)P_3_ inhibited the vertex enrichment of Ypt7 and the HOPS subunit Vps33. Instead of accumulating at vertices, both were distributed over the entirety of vacuole membranes. We did not perform a systematic check of all known vertex components; however, we know that eliminating any protein or lipid from vertices interferes with microdomain formation/stability and downstream events, including trans-SNARE complex formation (Karunakaran and Fratti, 2013; Collins and Wickner, 2007; Jun and Wickner, 2007).

### PI(3,4,5)P_3_ and SNARE function

As predicted from the vertex formation data, we found that blocking PI(3,4,5)P_3_ with Grp1 or modifying it with PTEN inhibited trans-SNARE complex formation. Interestingly, both Grp1 and PTEN triggered a disproportionate release of Vam7 from vacuoles. This was coupled with the uneven release of Vps33 and Vps18. More Vps33 was released compared to Vps18, suggesting that HOPS may disassemble without free PI(3,4,5)P_3_, which would not be due to direct binding, as HOPS has been shown not to bind the lipid (Stroupe et al., 2006). Regarding Vam7, it was unclear whether its release was due to direct interactions with PI(3,4,5)P_3_ or an indirect result of faulty vertex assembly that would interfere with Vam7 binding PI3P, SNAREs or HOPS.

The interaction between Vam7 and PI3P has long been recognized as the mechanism by which Vam7 associates with membranes, prior to interactions with its cognate SNAREs and HOPS (Cheever et al., 2001; Boeddinghaus et al., 2002). Its lipid binding capacity lies in its N-terminal PX domain that mediates PI3P binding, which can be largely abolished by mutating Tyr42 to Ala. That said, Vam7^Y42A^ can still associate with vacuoles and support fusion at reduced levels (Fratti and Wickner, 2007). By microscopy, Cheever et al. showed that GFP-Vam7^Y42A^ was largely shifted from the vacuole to the cytoplasm. While mostly hazy in the cytoplasm, a non-trivial amount of GFP-Vam7^Y42A^ seemed to localize to vacuole-adjacent puncta, which are likely pre-vacuolar compartments. The images were not quantified so it was hard to tell how much was shifted. Interestingly, the authors also showed that ts*vps34* at the non-permissive temperature only partially shifted Vam7 to the cytosol.

Previously, the low activity of Vam7^Y42A^ had been solely attributed to its interactions with proteins and not other lipids. However, recent work has shown that PX domains from other proteins can bind to alternative lipids through second binding sites (Chandra et al., 2019). Based on this, we tested if Vam7 could interact with PI(3,4,5)P_3_. Studies using the Vam7 PX domain alone and lipid overlay experiments have shown mixed results (Yu and Lemmon, 2001; Cheever et al., 2001). Using membraneless BLI experiments, in which biotinylated lipids were attached to streptavidin probes, we found that Vam7 did in fact bind to b-PI(3,4,5)P_3_ with a slightly higher affinity compared to b-PI3P. This was likely due to the additional contacts generating a stronger binding energy. Importantly, Vam7^Y42A^ failed to bind b-PI3P as expected. Yet, it was able to bind b-PI(3,4,5)P_3_, albeit with reduced affinity compared to WT Vam7. These data suggest that Vam7 uses a second binding site to interact with PI(3,4,5)P_3_; however, this does not rule out interactions through the PI3P binding pocket. Vam7^Y42A^-PI(3,4,5)P_3_ binding could be due to interactions with residues in the pocket independent of Tyr42.

To address the binding of PI(3,4,5)P_3_, we use two computational approaches. First, we performed co-folding predictions of full-length VAM7 to short chain PI(3,4,5)P_3_ and PI3P and verified enrichment of these models performing molecular docking of DAG, PA, PI, PI3P, and PI(3,4,5)P_3_. Interaction zones were drawn around the PX-lipid binding pocket to orient interactions and exclude outer PBR regions that could also interact with anionic lipids. Boltz-2 predictions indicated that PI(3,4,5)P_3_ could bind the PI3P pocket with a predicted binding affinity of 49 nM and excluded interactions with Tyr42. Instead, binding was dependent on Arg88 to bind the D5 position while the D3 and D4 positions were organized by Arg41 and Ser43.

The second approach used the solution NMR structure of PX in MD simulations with membranes. The MD simulations provided a membrane-level explanation for why Vam7 can display different apparent sensitivities to the Y42A mutation depending on the phosphoinositide ligand. Three observations are most relevant. First, WT Vam7 on PI3P membranes formed the most reproducible membrane-bound state, with stable binding in all four simulation replicas. Second, the Y42A mutation strongly reduces the recurrence of this state, consistent with the fact that Tyr42 contributes to the canonical PI3P-recognition geometry. Third, PI(3,4,5)P_3_-containing systems show broader engagement of Lys/Arg residues, indicating that the more phosphorylated PI(3,4,5)P_3_headgroup can stabilize the protein through a distributed electrostatic interaction network. This distinction is important because the simulations do not simply support that one lipid is better than the other for membrane binding. WT+PI3P resulted in the most reproducible (canonical) binding geometry. PI(3,4,5)P_3_, in contrast, engaged a broader range of Lys/Arg–PI oxygen contacts.

The Y42A+PI3P simulations further clarify the mechanism. The mutant still forms several Lys/Arg PI3P contacts, but they did not result in the reproducible WT-like PI3P-bound geometry. This observation suggests that electrostatic contacts alone are insufficient to organize the canonical PI3P-bound pose. Tyr42 appears to contribute to the geometry or stabilization of this canonical state, while direct headgroup engagement is dominated by Lys/Arg residues. Thus, the simulations distinguish loss of a canonical PI3P-binding geometry from complete loss of membrane electrostatic engagement.

Multiple basic residues can engage PI(3,4,5)P_3_ oxygen atoms on the membrane-facing surface of the protein. Because PI(3,4,5)P_3_ has additional phosphate groups relative to PI3P, it can support a larger set of salt-bridge-like ion-pair contacts. This provides a plausible explanation for why PI(3,4,5)P_3_-associated membrane engagement can persist even when the canonical Tyr42-sensitive PI3P-recognition mode is weakened, without requiring a single WT-like PI3P-bound pose.

Overall, the simulations support a model in which Vam7 PX-domain membrane recognition includes both a canonical PI3P-recognition component and a broader electrostatic membrane-retention component. WT+PI3P best reports the canonical binding geometry, whereas PI(3,4,5)P_3_-containing membranes reveal the capacity of the PX domain to engage a more extended Lys/Arg-rich surface. This model reconciles the depth, residue-geometry, contact, and snapshot analyses and provides a mechanistic basis for interpreting the different effects of Y42A on PI3P- and PI(3,4,5)P_3_-dependent membrane binding.

### What is the role for dual lipid interactions with Vam7?

Although Vam7 can bind to both PI3P and PI(3,4,5)P_3_, we know that it can be released from vacuoles when either lipid is blocked or modified in the presence of the other. This poses the question: Why would two seemingly redundant lipid interactions be preserved? PI3P is delivered to vacuoles via endolysosomal trafficking in low concentrations and the pool of PI3P is regenerated by a new round of production on site (Thorngren et al., 2004). It is possible that the ebb and flow of PI3P levels could negatively impact Vam7 recruitment to vacuoles and that a second ligand *i.e*., PI(3,4,5)P_3_, is needed for stable association with the membrane. Could a lipid exchange occur where PI(3,4,5)P_3_ initially binds while PI3P pools are replenished early in the cycle? Since we added Grp1-PH and PTEN at the start of the trans-SNARE pairing and Vam7 retention experiments, it could be that Vam7 was released from PI(3,4,5)P_3_ when PI3P levels were still low. Originally, we hypothesized that PA could be the initial binding partner and that its conversion to DAG by Pah1 would lead to a hand off to PI3P. This was based on the required conversion of PA to DAG during pre-priming and the idea that Vam7 only binds one lipid at a time (Sasser et al., 2012; Starr et al., 2019, 2016). Now we must consider that PI(3,4,5)P_3_ could be the binding initiator, and that Vam7 could bind two lipid species in sequence or simultaneously.

Another question to consider is whether Vam7-PI(3,4,5)P_3_ interactions affect Ypt7 and HOPS enrichment at vertex microdomains. This is not unlikely as both Vam7 and Ypt7 interact with the HOPS complex, although not necessarily at the same time as seen by pulldown experiments (Price et al., 2000a; Seals et al., 2000; Brett et al., 2008; Stroupe et al., 2006; Collins et al., 2005; Fratti and Wickner, 2007; Fratti et al., 2007). Thus, it could be that changes in Vam7 conformation induced by PI(3,4,5)P_3_ binding influences how HOPS interacts with Ypt7. A more direct effect would be through disrupting Vam7-Ypt7 binding. A yeast-two-hybrid screen in the absence of PI(3,4,5)P_3_ showed that Vam7 and Ypt7 interact directly (Uetz et al., 2000). In this scenario PI(3,4,5)P_3_ would disrupt Vam7 binding to Ypt7 to promote interactions with other SNAREs and HOPS, respectively. More experimentation is required here.

Finally, we showed that PI(3,4,5)P_3_ affects 3Q-SNARE complex formation. Our data showed that PTEN and Grp1-PH led to the exclusion of Vti1 and Vam7 from CBP-Vam3 complexes. The absence of Vam7 is attributed to its release from the membrane, but Vti1 has a TMD and was not displaced from vacuoles. Interestingly, the hydrophilic helical region adjacent to the Vti1 TMD has a PBR with multiple Lys and Arg residues that could participate in binding negatively charged lipids including PI(3,4,5)P_3_, however, this remains to be tested. Other SNAREs including Syntaxin-1 and Syntaxin-17 have TMD-adjacent PBRs that bind to PIs to promote function in chromaffin granule secretion and autophagy, respectively (Laczkó-Dobos et al., 2024; Lam et al., 2008). Future studies will test whether the Vti1 PBR promotes binding to Vam3 followed by Vam7, if Vam7 binds Vam3 before Vti1, or if complex formation is concurrent. Alternatively, Vam7-membrane interactions could be needed for it to bind Vti1 and Vam3 in a 3Q-SNARE complex. We must note that a previous study showed that Vti1-Vam3 binding is not affected by eliminating or blocking other PIs, including PI3P, PI4P, and PI(4,5)P_2_ (Collins and Wickner, 2007). Also, in the absence of TMDs, the truncated Vti1 and Vam3 easily form soluble 3Q complexes with Vam7 in the absence of membranes (Jun et al., 2006). This underscores the importance of PI(3,4,5)P_3_ in SNARE function.

In conclusion, this study has shown that PI(3,4,5)P_3_ is critical in the regulation of vacuole homotypic fusion at multiple steps. While this study ends at the trans-SNARE complex stage, the gain of resistance data suggest that additional down-stream stages could be affected by PI(3,4,5)P_3_. Beyond vacuole fusion, it is likely that PI(3,4,5)P_3_ signals through the yeast orthologs of PDK1 (Pkh1/2), AKT1 (Sch9), PKC (Pkc1) and MAPK (Fus1, Kss1, Mpk1) to influence other functions such as autophagy, Ca^2+^ transport, and actin remodeling (Brady et al., 2006; Inagaki et al., 1999; Ni et al., 2013; Levina et al., 2022; Asano et al., 2008; Dieterle et al., 2014; Kassouf et al., 2015; Hsu et al., 2000).

## Materials and methods

### Reagents

Chemical reagents were solubilized in PIPES-Sorbitol (PS) buffer (20 mM PIPES-KOH, pH 6.8, 200 mM sorbitol) with 125 mM KCl unless indicated otherwise. PIPES [Piperazine-N-N’-bis(2-ethanesulfonic acid)], NEM (N-ethylmaleimide), Coenzyme A (CoA), Creatine kinase, and reduced glutathione were purchased from Sigma-Aldrich (St. Louis, MO) and dissolved in PS buffer or DMSO. FM4-64, Goat anti-rabbit IgG (H+L) secondary antibody DyLight 650 conjugate, Fluoromount-G with DAPI, Pierce glutathione agarose, and Phosphate Sensor were from Thermo-Fisher (Waltham, MA). Chitin resin was from New England Biolabs (Ipswich, MA). Fluorescent CF488 goat-anti GST was from Biotium (Fremont, CA). Poly-L-Lysine was from Santa Cruz Biotechnology (Dallas, TX). Creatine phosphate was from Abcam (Waltham, MA). PIT-1, 3,5-dimethyl PIT-1 (DMPIT), and SAR405 were from Cayman Chemical (Ann Arbor, MI) and dissolved in DMSO. 3-nitrocoumarin (3NC) was from Med Chem Express and dissolved in DMSO (Monmouth Junction, NJ). C8-PC (1,2-dioctanoyl-sn-glycero-3-phosphatidylcholine), C8-PE (phosphatidylethanolamine), and C8-PS (phosphatidylserine) were from Avanti Polar Lipids (Alabaster, AL). C8-PI3P (1,2-dioctanoyl-sn-glycero-3-phosphatidylinositol 3-phosphate), C8-PI(3,5)P_2_, C8-PI(4,5)P_2_, C8-PI(3,4,5)P_3_, biotin-PI3P (b-PI3P), b-PI(3,4)P_2_, b-PI(3,5)P_2_, b-PI(4,5)P_2_, b-PI(3,4,5)P_3_, Inositol-1,3,4-trisphosphate (Ins(1,3,4)P_3_), Ins(1,3,5)P_3_, Ins(1,4,5)P_3_, Ins(1,3,4,5)P_4_, PI(3,4,5)P_3_ mass ELISA kit, and PI(4,5)P_2_ mass ELISA kit were from Echelon Biosciences (Salt Lake City UT). *p*-nitrophenyl phosphate was from MP Biomedicals (Santa Ana, Ca). Calmodulin agarose was Agilent (Santa Clara, CA). Octet Streptavidin (SA) biosensors were from Sartorius (Göttingen, Germany). PreScission Protease was from GenScript (Piscataway, NJ).

### Plasmids and Recombinant proteins

Recombinant GST-Vam7, GST-Vam7^Y42A^, GDI, Pbi2 (Inhibitor of proteinase B), GST-ENTH and Oxalyticase were prepared as described previously (Fratti and Wickner, 2007; Miner et al., 2016; Fratti et al., 2007; Starai et al., 2007; Slusarewicz et al., 1997; Scott and Schekman, 1980). Plasmids to produce GST-Grp1-PH and GST-PTEN were from Addgene (Watertown, MA). The plasmid to make GST-Grp1-PH^K273A^ was a gift from Dr. N. Leslie (Heriot Watt University, Edinburgh, UK) (Lindsay et al., 2006). Plasmids were transformed into *E. coli* BL21 DE3 pLysS (New England Biolabs) and induced with 0.1 µM or 0.1 mM IPTG for GST-PTEN and GST-GRP1 constructs, respectively, and incubated for 16-18 h at 18°C. GST-tagged proteins were isolated using standard methods with glutathione agarose, eluted with reduced glutathione and dialyzed against PS buffer with 125 mM KCl. The GST tag was removed from GST-ENTH with PreScission protease following the manufacturer’s instructions then dialyzed against PS buffer with 125 mM KCl.

### mCherry-Grp1-PH yeast plasmid

A DNA fragment encoding GST-linker-mCherry-Grp1-PH with 5’-BamHI and 3’-HindIII restriction sites was synthesized by Twist Bioscience (South San Francisco, CA). This was amplified by PCR using forward primer GST-mCherry-PH TTCTGTAAGCGCggatcc and reverse primer GST-mCherry-PH GTGGCTATCCCCaagcttc containing BamHI and HindIII restriction sites (lowercase). The amplicon and the shuttle vector p405ADH1(Addgene) were digested with BamHI-HF and HindIII-HF restriction enzymes and ligated into the vector so expression was under the strong constitutive *ADH1* promoter. The ligation product was transformed into chemically competent DH5α cells, and transformants were selected and sequenced to confirm the correct insertion of the construct. Linearized plasmid was transformed into yeast by standard lithium acetate methods and selected for growth using synthetic media without Leu.

### Strains, Vacuole isolation and fusion

Yeast strains **(Table S3)** were grown in YPD (1% yeast extract, 2% peptone, 2% dextrose) or synthetic drop-out media without Trp, Ura or Leu. The pH of drop out media was adjusted to 6.0. The temperature sensitive *VPS34* mutant (ts*vps34*) was a gift from Dr. D. Klionsky (University of Michigan) (Wurmser and Emr, 1998). The temperature sensitive *MSS4* mutant (ts*mss4*) was from Dr. David Drubin (University of California at Berkeley) and Dr. Aaron Neiman (Stony Brook University) (Audhya and Emr, 2003; Nunez et al., 2023). Vacuoles were isolated as described (Haas et al., 1994). In vitro fusion reactions (30 µl) contained 3 µg each of vacuoles from BJ3505 (*PHO8 pep4*Δ) and DKY6281 (*pho8*Δ *PEP4*) backgrounds, reaction buffer (PS buffer, 125 mM KCl, 5 mM MgCl_2_), ATP regenerating system (1 mM ATP, 0.1 mg/ml creatine kinase, 29 mM creatine phosphate), 10 µM CoA, and 283 nM Pbi2 (Protease B inhibitor) (Jones et al., 1982; Klionsky and Emr, 1989). Fusion was determined by the processing of pro-Pho8 (alkaline phosphatase) from BJ3505 by the Pep4 protease from DKY6281. Fusion reactions were incubated at 27°C for 90 min and Pho8 activity was measured in developer solution (250 mM Tris-HCl pH 8.5, 0.4% Triton X-100, 10 mM MgCl_2_, and 1 mM *p*-nitrophenyl phosphate). Pho8 activity was stopped after 5 min by addition of 1 M glycine pH 11 and fusion units were measured by determining the *p-*nitrophenolate produced by absorbance at 400 nm. One unit of fusion is 1 µmol *p-*nitrophenolate produced per min per µg BJ3505 (*PHO8*) vacuoles (Haas et al., 1994; Conradt et al., 1994). Due to the variability in daily vacuole preparations the control maximum fusion can range by multiple U. Thus, we normalize each experiment to its own control maximum set to 1.

### Trans-SNARE complex isolation

Trans-SNARE pairing was measured as previously described with some modifications (Collins and Wickner, 2007; Jun et al., 2007; Qiu and Fratti, 2010; Sasser et al., 2012, 2013). Large scale 15X (450 µL) fusion reactions containing 45 µg of DKY6281 vacuoles (*VAM3 NYV1*) and 45 µg of BJ3505 vacuoles that lacked *NYV1* and contained Vam3 containing an internal calmodulin binding peptide (CBP) between the H_abc_ and SNARE domains (*CBP-VAM3 nyv1*Δ) (Collins and Wickner, 2007). Reactions were treated with buffer alone or 2 mM NEM as a negative control to prevent SNARE activation. Separate reactions were treated with GST-PTEN or GST-Grp1-PH at the indicated concentrations. Reactions were incubated for 60 min at 27°C, then placed on ice for 5 min before centrifugation (13,000 g, 15 min, 4°C) to pellet membranes. The supernatants were carefully removed, and the pellets were overlaid and resuspended with 200 µL cold solubilization buffer (SB: 20 mM Tris-Cl, pH 7.5, 150 mM NaCl, 1 mM MgCl_2_, 0.5% Nonidet P-40 alternative, 10% glycerol) with protease inhibitors (1 mM PMSF and 1 protease inhibitor tablet per 10 ml SB). Reactions were brought up to 600 µL with additional SB and nutated at 4°C for 20 min. Insoluble debris was removed by centrifugation (16,000 g, 20 min, 4°C). Solubilized (580 µL) material was transferred to pre-chilled tubes and 58 µL were removed from each reaction as 10% of the total reactions. CaCl_2_ was added to each sample to a final concentration of 2 mM. Next, 50 µL of calmodulin beads equilibrated with SB was added to each reaction. Mixtures were nutated overnight at 4°C. CBP-Vam3 complexes bound material was collected by centrifugation (1,300 x g, 2 min, 4°C) and washed 4 times with 600 µL fresh ice-cold SB. Calmodulin bound material was eluted with 1X SDS loading buffer containing 5 mM EGTA and boiled for 5 min. Samples were resolved by SDS-PAGE, transferred to nitrocellulose and probed with antibodies against Vam3, Vam7, Nyv1, Vti1, Vps33 and Vps18. Bound primary antibodies were visualized with DyLight 650-Goat anti-rabbit IgG (H+L).

### Fluorescence microscopy of vacuoles and vertex microdomain formation

Isolated vacuoles from BJ3505 or from cells expressing GFP-Ypt7 or Vps33-GFP were subjected to docking assays as previously described (Fratti et al., 2004; Wang et al., 2002) with slight modifications. Reactions (30 μL) contained 6 μg of vacuoles isolated from the indicated strains in fusion reaction buffer modified for docking conditions (PS buffer, 100 mM KCl, 0.5 mM MgCl_2_, 0.33 mM ATP, 13 mM creatine phosphate, 33 μg/mL creatine kinase, 10 µM coenzyme A, and 280 nM IB_2_). Measuring the vertex enrichment of factors during tethering and docking was performed with vacuoles from cells expressing GFP fusion proteins or labeled with lipid binding probes. To track GFP-Ypt7 and Vps33 localization reactions were incubated under docking conditions as described above and stained with 4 μM FM4-64 prior to examination (Wang et al., 2003; Fratti et al., 2004). To localize the distribution of PI(3,4,5)P_3_ on vacuoles, reactions were incubated with 150 nM GST-Grp1-PH. GST was then visualized with fluorescent (CF488) goat-anti-GST antibody (1:5000). Briefly, reactions were treated with PS buffer, DMSO, PTEN or SAR405 for 5 min, followed by adding 150 nM GST-Grp1-PH and incubating for 5 min. Next, CF488-anti-GST antibody was added to each reaction and further incubated for 20 min. Following incubation, reactions were gently centrifuged to remove unbound material, resuspended with PS buffer and mixed with 20 μL of 0.6% low melt agarose in PS buffer melted at 50°C and cooled prior to mixing with vacuoles. Next, 20 μL aliquots were mounted on pre-chilled slides and observed by fluorescence microscopy. Images were acquired using a Zeiss Axio Observer Z1 inverted microscope equipped with an X-Cite 120XL light source, Plan Apochromat 63X oil objective (NA 1.4), and an AxioCam CCD camera. CF488 was visualized using a 38 HE EGFP shift-free filter set and FM4-64 was visualized with a 42 HE CY3 shift-free filter set. Exposure times were set using WT vacuoles for each fluorescence channel and scripts acquired non-specific images followed by specific reporters. This ensured that bleaching was consistent to negate it as a factor in calculating intensity ratios. Exposure times were held constant within an experiment.

Images were analyzed using ImageJ software (NIH). Vertex enrichment was determined by first measuring maximum fluorescence intensity in each channel at each contact point between membranes, *i.e.*, vertex domain within a cluster. Next, fluorescence intensity was measured in each channel at outer membrane domains where vacuoles are not in contact with other membranes. The ratio of specific (*e.g.,* GFP) to non-specific (FM4-64) was determined for vertices and outer membrane domains and compared for relative enrichment. Measurements for each condition were taken of 15-20 clusters to yield 100-300 vertices for each condition/strain per experiment. Data from multiple experiments were combined cumulative it distribution plots and scatter column plots showing individual values as well as the geometric means and geometric standard deviation for each condition.

### Fluorescence microscopy of whole cells

Cells were inoculated in YPD or complete synthetic medium without Ura or Leu and grown overnight at 30°C. Cultures were diluted to 0.25 OD_600_ into 10 ml of fresh YPD incubated until OD_600_ reached 0.4-0.6 at 30°C or 37°C. Cells were fixed with 3.7% Formaldehyde for 60 min. Fixed cell pellets were collected and washed twice with 1 ml 1X PBS buffer and resuspended in 0.5 ml spheroplast buffer (50 mM KPi buffer pH 7.5, 600 mM Sorbitol, 0.2% YPD). Cell suspensions (200 µL) were mixed with 70 µg oxalyticase and 0.5 µL stock 2-mercaptoethanol and incubated in a 30°C water bath for 60-90 min. Cells were then washed once with 1 ml of 1X PBS + 0.05% Tween 20 and resuspended in 50 µl of PBS + 0.05% Tween 20. Then 40 µl of cells were transferred onto coverslips coated with 0.1% Poly-L-lysine in a 6-well dish and incubated on a shaker at room temperature for 5-10 minutes. After incubation, the fluid was aspirated and coverslips were washed 3 times with 1 mL 1X PBS buffer to ensure that they remained submerged during each wash. Coverslips were dried and cells were blocked with PBS + 1mg/ml BSA for 30 min at room temperature, then washed 3 times with 1X PBS buffer. Next, a 10 µM GST-Grp1-PH solution diluted with PBS with 1mg/ml BSA was added to coverslips and incubated for 1 h at room temperature. After incubation, the coverslips were washed 3 times with 1X PBS buffer and incubated with 1:5000 CF-488 anti-GST fluorescent antibody diluted in PBS + 1mg/ml BSA in the dark for 30 min. Coverslips are washed 3 more times with 1X PBS buffer and are dried for a few minutes before mounting onto pre-cleaned microscope slides with 10 µl mounting media (fluoromount-G^TM^ with DAPI) followed by imaging by fluorescence microscopy.

Images were analyzed using ImageJ software (NIH). Maximum fluorescence intensities were measured using a DAPI filter or a GFP filter set for CF488. The ratio of CF488-Grp1-PH (PI(3,4,5)P_3_) to DAPI was determined for each cell. Measurements for each condition were taken of ∼100 cells per condition per experiment. Data from multiple experiments were combined in column plots showing individual values as well as the geometric means and geometric standard deviation for each condition.

For live cells, well-focused cells were analyzed using ImageJ (NIH). Regions of interest (ROIs) were manually defined for each cell, and the Area, Raw Integrated Density, and Mean Gray Value were measured for the mCherry and MDY-64 fluorescence channels. Background fluorescence was measured from a cell-free region within the same field of view. Corrected Total Cell Fluorescence (CTCF) was calculated for each channel using the following equation: CTCF = Raw Integrated Density − (Area × Background Mean Gray Value). The mCherry CTCF of each cell was normalized to the corresponding MDY-64 CTCF by calculating the ratio of mCherry CTCF to MDY-64 CTCF.

### Immunoblotting

Vacuoles were solubilized with 95°C Laemmli buffer for 5 min. Extracts were resolved using 10% SDS-PAGE and transferred to nitrocellulose for immunoblotting. Rabbit antibodies against Actin, Nyv1, Vam3, Vps18, Vps33 and Ypt7 were prepared as described (Eitzen et al., 2002; Nichols et al., 1997; Seals et al., 2000; Haas et al., 1995). Goat anti-rabbit IgG (H+L) antibody DyLight 650 conjugate was used as a secondary antibody. Fluorescence was measured with an Azure 400 gel documentation system (Azure Biosystems, Dublin, CA).

### Bio-Layer Interferometry (BLI)

Binding to lipids was measured by BLI as described (Calderin et al., 2025). Biotinylated phosphoinositides were resuspended in PS buffer to a final stock concentration of 0.1 mM. Lipids were diluted to 500 nM with BLI running buffer (PBS with 0.002% Tween-20, v/v) and 190 µL were added to wells in a 96-well microplate. GST-Vam7, and GST-Vam7^Y42A^, GST-Grp1-PH, GST-Grp1-PH^K273A^ were diluted with BLI running buffer and 190 µL of each dilution, for each analyte, was loaded to corresponding wells. Binding curves were generated in Graphpad Prism and K_D_ values were calculated using non-linear regression and one-site specific binding.

### Lipid extraction of acidic PIs for mass ELISA

Yeast vacuoles were isolated as stated above and prepared for docking conditions as used for microscopy experiments. Reactions were incubated for 30 min at 27°C. For extraction of lipids from temperature sensitive strains (ts*vps34* and ts*mss4*), reactions were incubated at 30°C or 37°C. After incubation, lipids were extracted as described with some modification (Hsu et al., 2000). First, 300 µL of chilled acetone was added to each reaction on ice for 5 min followed by centrifugation (3,000 g, 5 min at 4°C). The supernatant was carefully removed (leaving a minimal amount to avoid disturbing the pellet), and 700 µL MeOH/CHCl_3_ (1:1) was added to resuspend the pellet, which was followed by vortexing heavily at RT for 10 min, or using an Eppendorf MixMate at 1850 RPM or 10 min. The material was centrifuged (3,000 g, 5 min at 4°C) and the supernatants were carefully discarded. The remaining pellets were resuspended with 700 µL MeOH:CHCl_3_:HCl (80:40:1) and mixed at RT for 45 min with a vortex or MixMate (at 1850-2000 RPM) until the pellets were dispersed in solution. Suspensions were transferred to new 15 mL conical tubes after which 750 µL CHCl_3_ and 1.35 mL 0.1 M HCl were added to each tube and vortexed for 5-10 min. Extractions were transferred to a new set of 15 mL conical tubes prior and centrifuged (3,000 g, 5-10 min at 4°C). Only 800 µL of the lower phases (∼1.5 mL total) were carefully transferred to 1.5 mL microcentrifuge tubes, dried in a speed vac or under a N_2_ stream and stored at -20°C until further use.

For whole cell analysis, strains were grown to OD_600_ ∼0.7 and prepared for spheroplasting treatment as mentioned above for yeast vacuole isolation. Fractions (200 µL) were collected from 1 L of resuspended cells in 15 mL spheroplast buffer, mixed with oxalyticase and incubated for 35 min. Following incubation, the cells were pelleted (3,000 g, 5-10 min, 4°C). Supernatants were removed, and the corresponding cell pellets were resuspended in 100 µL of PS buffer and added to 3X fusion conditions as mentioned above and incubated for 90 min. Lipids were then extracted following the same protocol for vacuoles.

### ELISA and Data Analysis

The ELISA Data for PI(3,4,5)P_3_ and PI(4,5)P_2_ were run and analyzed according to Echelon Biosciences Inc., as shown elsewhere (Gross et al., 2015; Costa et al., 2015). Lipid films were resuspended in Kit-specified buffer and set on Eppendorf MixMate at 1850 RPM for 20 min. Each trial, a distinct extraction from a distinct overnight growth, and standard curves were analyzed with GraphPad Prism using a Sigmoidal dose response-variable slope curve (four-parameter logistic, 4PL) analysis. This was performed through choosing the “RIA or ELISA - interpolate unknown from Sigmoidal Curve” on GraphPad Prism then converting the standard curve values for standard curve value to log(concentration). The data presented was in log(concentration), hence taking 10^(log(concentration), converted the data back into concentration (pmol). If a specific trial had a dilution factor, then adjusted by either multiplying or dividing by the factor. Then among similar conditions, the results are averaged together and compared.

### Phosphatase activity

Phosphatase activity was measured using the Phosphate Sensor (Invitrogen) according to the manufacturer’s instructions with modifications. Enzymatic reactions were performed in reaction buffer containing 50 mM Tris-HCl (pH 7.6), 25 mM NaCl, and 0.5 mM DTT (added fresh). Solutions were made with Molecular Biology Grade (Corning) water to lower background fluorescence. PTEN (727 nM) was incubated with 120 µM C8-PI(3,4,5)P_3_ in a total volume of 10 µL in black, non-binding surface 384-well microplates (Corning). Phosphate Sensor (diluted to ∼0.001 pmol per well) in detection buffer (20 mM Tris-HCl, pH 7.6) was added. Fluorescence was measured using a microplate reader with excitation and emission wavelengths of 430 nm and 450 nm, respectively. Activity was calculated by subtracting background fluorescence from controls and interpolating corrected relative fluorescence values.

### Computational modeling

Amino acid sequences were procured from the SGD for *VAM7* (YGL212W) (Cherry et al., 2012). Short chain lipids were prepared in Schrödinger Maestro and LigPrep (Schrödinger Inc, New York, NY). Structures were exported using .sdf format and amino acid sequences were run with short chain lipids with Boltz2 (Passaro et al., 2025) using standard settings with no residue modifications, pocket restraints, or covalent restraints with MSA mode mmseqs2_uniref_env, number recycles 6, sampling steps 200, diffusion samples 5, and step scale 1.5. Complex pLDDT (predicted local distance difference test) for Boltz-2 predictions are: Vam7 alone .6502, Vam7 to PI3P .6988, and Vam7 to PI(3,4,5)P_3_ .6628. Predicted IC_50_ (-logIC_50_) values are transformed (kcal/mol = (6 – log(IC_50_)) × 1.364) to predicted binding affinity for models using short-chain lipids as reported in **Fig.11C**.

Top ranked poses for each *de novo* co-folding prediction from Boltz-2 were then exported in .cif format and uploaded into Schrödinger Maestro. Protein Preparation Wizard Schrödinger Release 2026-1: Protein Preparation Workflow; Epik, Impact, and Prime in Schrödinger were run using OPLS4 forcefield OPLS4 (Lu et al., 2021). Receptor grid generation using short-chain lipids of interest was used as the center of 35x15 Å grid box using the Schrödinger grid generation tool. Glide SP docking was performed for g-scores reported in **Fig.11C** using enhanced and expanded sampling. Glide XP was performed for g-scores reported in **Fig.11D** with 5 pose output, keeping the top 3 outputs for each short-chain lipid (Friesner et al., 2004). G-scores are reported with respect to the top 3 inputs for each short-chain lipid of interest. Ligand Interaction Diagrams were exported for the Boltz-2 predictions after minimization with OPLS4.

### Molecular dynamics

Molecular dynamics (MD) simulations were performed to compare binding of WT and Y42A Vam7 PX domains to membranes containing either PI3P or PI(3,4,5)P_3_. The starting structure was the Vam7 PX domain obtained from PDB 1KMD (Cheever et al., 2001; Lu et al., 2002). The simulated construct corresponded to the isolated PX domain, with ACE and CT3 caps added to the N- and C-termini, respectively. The Y42A mutant was generated from the same PX-domain model by replacing Y42 with an alanine.

Four protein–membrane conditions were simulated: WT+PI3P, WT+PIP3, Y42A+PI3P, and Y42A+PIP3. Each condition included 4 independent replicas, resulting in 16 production simulation runs in total. Membranes contained POPC, POPE, POPS, ergosterol, and either PI3P or PI(3,4,5)P_3_ at a molar ratio of 50:20:10:10:10. Initial bilayers were generated with CHARMM-GUI (Jo et al., 2008). and converted to a highly mobile membrane mimetic (HMMM) representation to accelerate sampling (Ohkubo et al., 2012; Hasdemir et al., 2026). Two independent membrane builds were used for each phosphoinositide composition, and different initial PX-domain orientations were used to reduce bias caused by the protein’s starting configuration relative to the membrane.

All systems were simulated with NAMD 3 (Phillips et al., 2005, 2020) using the CHARMM36m protein force field (Huang et al., 2017) and CHARMM36-compatible lipid parameters (Klauda et al., 2010). Systems were solvated with TIP3P water (Jorgensen et al., 1983) and ionized with KCl. Long-range electrostatics were treated with particle-mesh Ewald (Darden et al., 1993). A 12-Å nonbonded cutoff was used, bonds involving hydrogen atoms were constrained, and production simulations were run with a 2-fs timestep. Each replica was equilibrated in two restrained stages, followed by 250 ns of production sampling. The full simulation set therefore contained 4.0 *µ*s of aggregate production sampling. Production trajectories were wrapped relative to the membrane, and water and ions were removed before analysis.

Trajectory analysis was performed using VMD Tcl and Python workflows (Humphrey et al., 1996). C_α_ RMSD was calculated after fitting each frame to the first frame using C_α_ atoms and was used only as a structural quality-control metric. Membrane binding was quantified using a depth metric defined as the lowest atom in the PX domain with respect to the membrane phosphate plane. A frame was considered stably membrane-bound when the depth of binding was ≤ 0 Å, and continuous membrane-binding lasted at least 30 ns. For stably bound frames, per-residue distances to the membrane phosphate plane were calculated to describe residue-level binding. Stably-bound per-residue distance profiles were dimensionally reduced with UMAP and clustered with DBSCAN to visualize conformational heterogeneity (McInnes et al., 2018; Ester et al., 1996). To quantify electrostatic phosphoinositide engagement, contacts were calculated between Lys/Arg side-chain nitrogen atoms and PI3P or PIP3 oxygen atoms using a 4.0-Å cutoff. Complete simulation details, equations, system tables, and analysis definitions are provided in the Supplementary Methods.

### Quantification and statistical analysis

Fusion results were expressed as the mean ± SE or geometric mean ± geometric SD as needed. Experimental replicates (n) are defined as the number of separate experiments. For comparison of vertex enrichment, all the ratio data was log-transformed to yield near-normal distribution with comparable variances. Non-parametric analysis gave indistinguishable results. Statistical analysis was performed by unpaired two-tailed t-test or One-Way ANOVA for multiple comparisons using Prism 10 (GraphPad, San Diego, CA). Statistical significance is represented as follows: \**p*<0.05, \*\**p*<0.01, \*\*\**p*<0.001, \*\*\*\**p*<0.0001. Tukey, Dunnett, and Šidák post hoc tests were used for multiple comparisons and individual p-values.

### Online supplemental material

**Fig. S1** related to **Fig. 8** shows experiments labeling of vacuoles with ENTH domain to visualize PI(4,5)P_2_. WT vacuoles were incubated at 30°C and 37°C to show that labeling was not directly sensitive to temperature changes. **Fig. S2** relates to the discussion showing Grp1-PH to visualized PI(3,4,5)P_3_. The experiment tested if blocking Fab1 activity with Apilimod affected PI(3,4,5)P_3_ production and labeling. **Fig. S3** is related to Fig. 13 and shows RMSD replicas for WT and Y42A interactions with PI3P or PI(3,4,5)P_3_. **Fig. S4** relates to Fig. 13 and shows the UMAP and the number of basic residue contacts of WT and Y42A interactions with PI3P or PI(3,4,5)P_3_. It also shows the distance of individual amino acids to the membrane phosphate layer.

## Data availability

All data generated or analyzed during this study are available upon request. Additional data sharing information is not applicable to this study.

## Acknowledgements

The authors wish to thank Dr. William Wickner for the generous gifts of antibodies, Dr. Nicholas Leslie for donated plasmids, and Drs. Daniel Klionsky, David Drubin, Aaron Neiman and Scott Emr for strains. We also thank Dr. Nick Wu and Timothy Tan for assistance with BLI assays. We also acknowledge Elliel Foley, Lia George, and Jordan Alioto for technical assistance. This research was supported by a grant from the National Science Foundation (MCB 2216742) to RAF, and the National Institutes of Health (R24-GM145965 to E.T.). JDC was partially supported by an NIGMS-NIH Chemistry-Biology Interface Training Grant (5T32-GM070421). We also thank Dr. Jennifer Grant for critical reading, editing and discussion of the manuscript.

## Author contributions

Conceptualization: C.Z., J.D.C., R.P.S., S.A., E.T. and R.A.F.; Data curation: C.Z., J.D.C., R.P.S, S.A., L.G., E.T. and R.A.F.; Formal analysis: C.Z., J.D.C. R.P.S., S.A., L.G., E.T. and R.A.F.; Investigation: C.Z., J.D.C., R.P.S., S.A., L.G., A.T., V.F., V.S., K.T.P. J.M.K., C.K., R.A., D.G., E.T. and R.A.F.; Methodology: C.Z., J.D.C., S.A., L.G., E.T. and R.A.F.; Visualization: S.A., L.G., E.T., R.A.F.; Writing-original draft: C.Z., J.D.C., S.A., E.T. and R.A.F.; Writing-review and editing: All authors; Resources: E.T. and R.A.F.; Supervision: E.T. and R.A.F.; Project administration: R.A.F.; Funding acquisition: E.T. and R.A.F.

---

**Figure S1.**
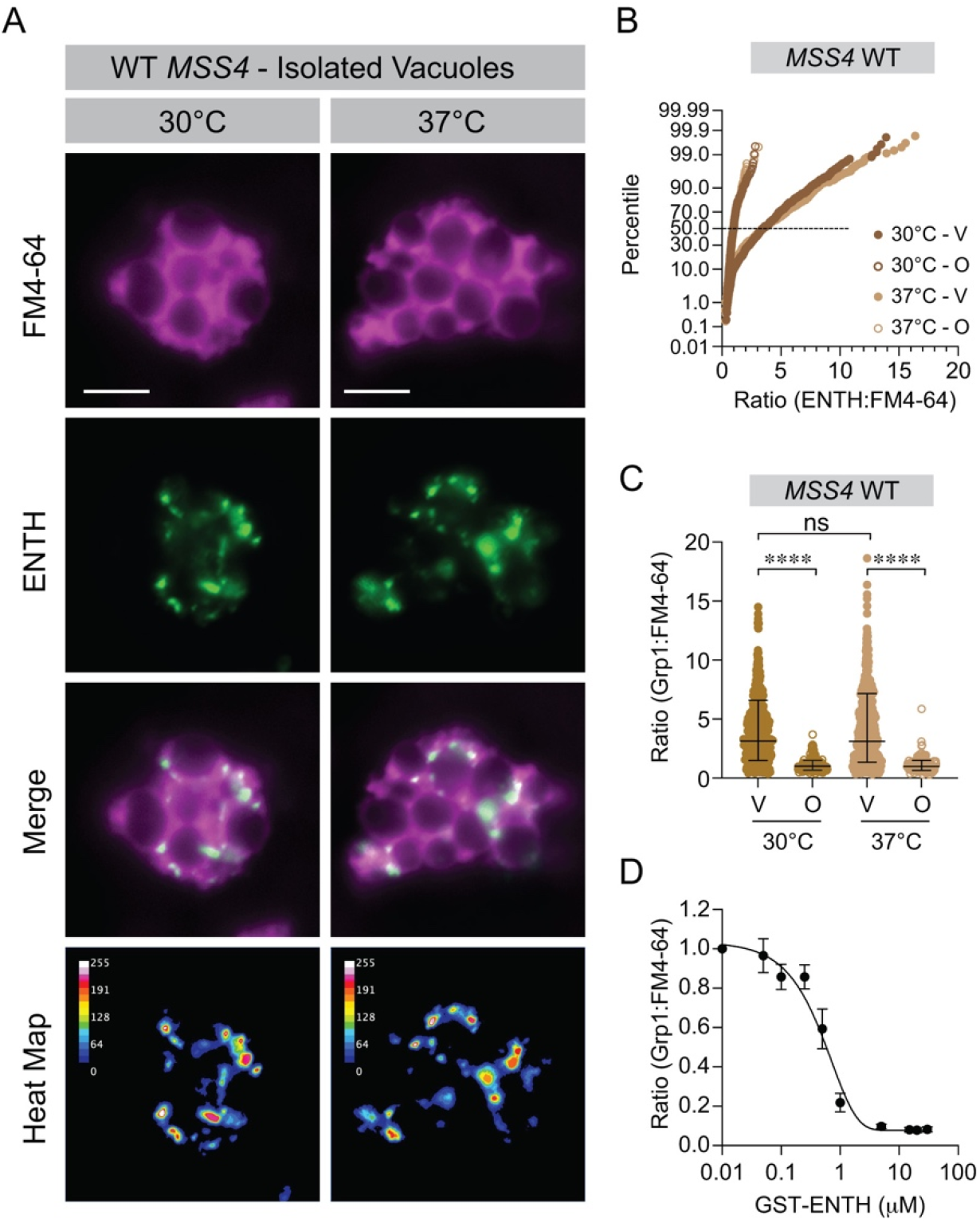
ENTH is not temperature sensitive with WT*MSS4*. **(A)** Docking reactions using purified WT *MSS4* vacuoles were incubated for 30 min at 30°C or 37°C and labeled with GST-ENTH as described. After incubation the reactions were place on ice and mixed with 5 µM FM4-64 to label entire vacuoles. Vacuoles were pelleted to remove excess GST-ENTH and antibody and resuspended in PS buffer. Reactions were next mixed with low melt agarose, mounted on glass slides and observed by fluorescence microcopy. Scale bars: 5 µm. **(B)** Images were analyzed and plotted in cumulative distribution plots as above (30°C-V = 495; 30°C-O = 207; 37°C-V = 568; 37°C-O = 196). **(C)** Scatter plots of pooled data shown in **(B)**. Error bars represent geometric means ± geometric standard deviations. Significance was determined using one-way ANOVA for multiple comparisons and Tukey’s test for individual *p*-values. \*\*\*\**p*<0.0001, ns – not significant. **(D)** Fusion inhibition by GST-ENTH. Results were normalized to the untreated control set to 1. Data represent mean ± SE (N=6).

**Figure S2:**
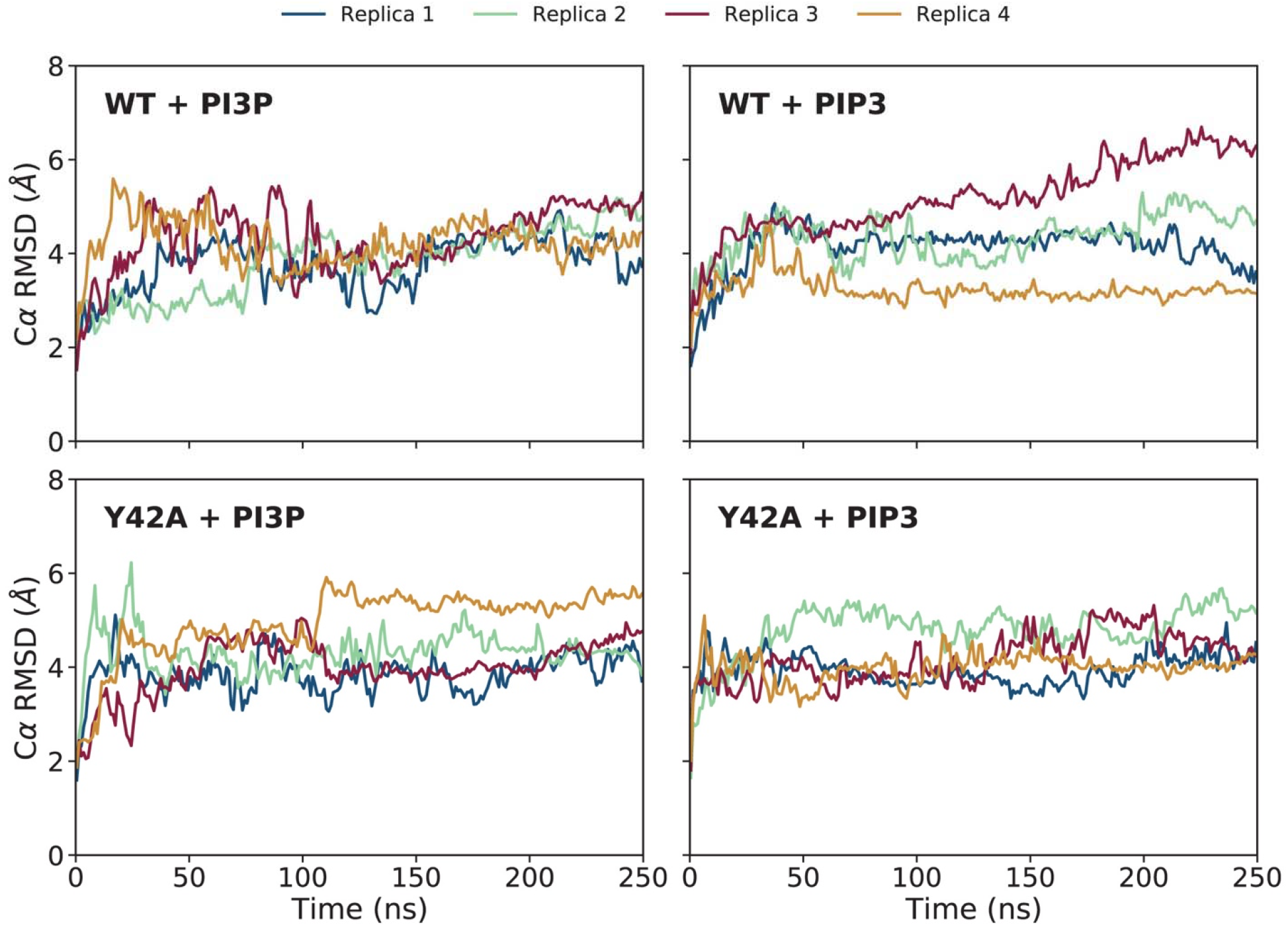
C_α_ RMSD for Vam7 PX domain simulations. C_α_ RMSD time series are shown for the 4 replicas of each simulated condition. RMSD was calculated after fitting each trajectory to the first frame using C_α_ atoms. For visualization, every 10 consecutive frames were block-averaged into one data point.

**Figure S3:**
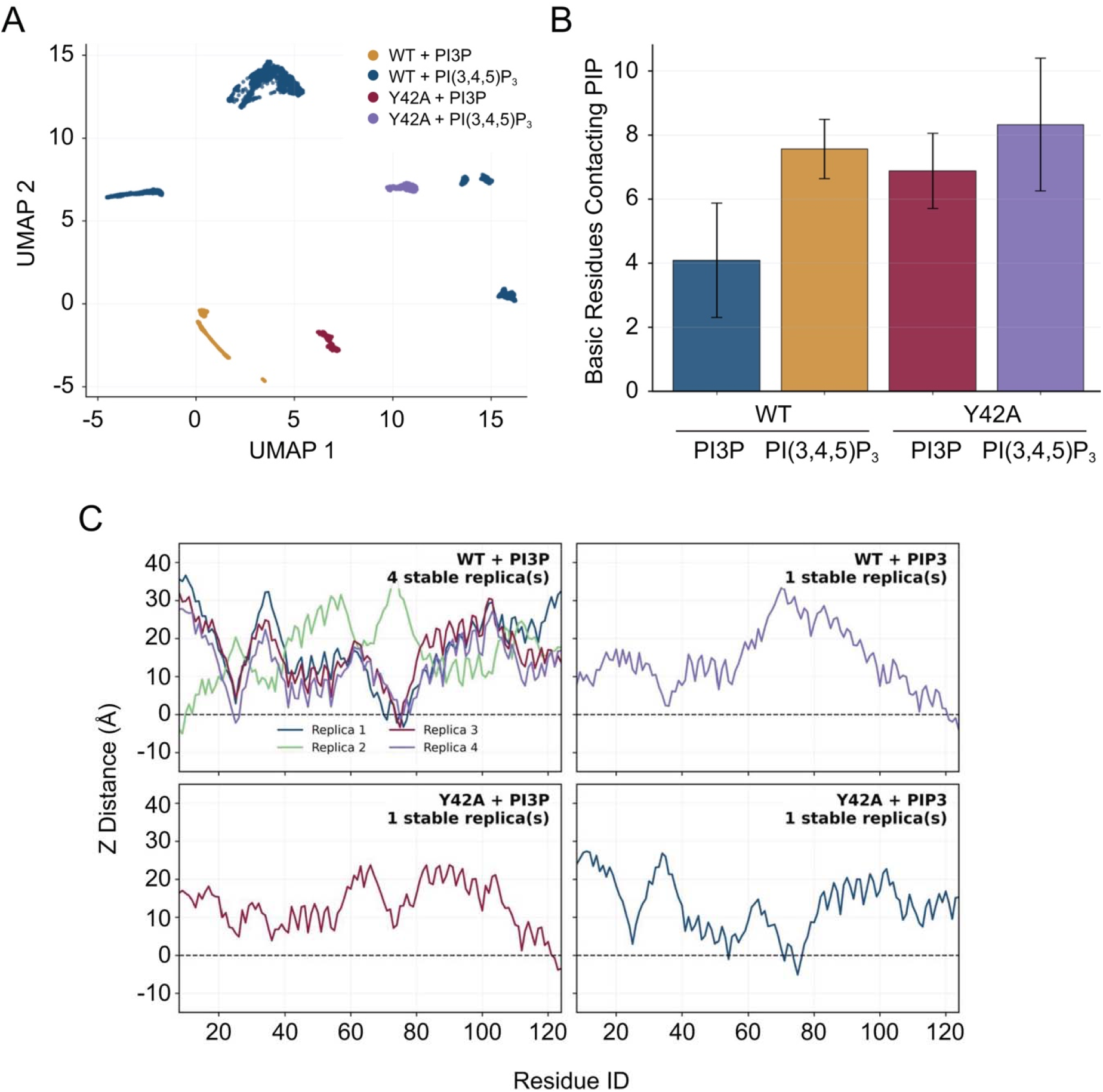
Conformational heterogeneity of stably bound forms. **(A)** UMAP projection of stably bound frames. WT–PI3P contributes the most to the largest stably bound ensemble, consistent with membrane binding in all 4 simulation replicas. **(B)** Mean number of Lys/Arg residues contacting PI3P or PI(3,4,5)P_3_ (PIP3) per frame across all production frames. Error bars indicate the standard deviations. **(C)** Per-residue membrane-distance profiles calculated from stably bound frames. WT–PI3P shows the most reproducible residue-level binding form.

**Figure S4.**
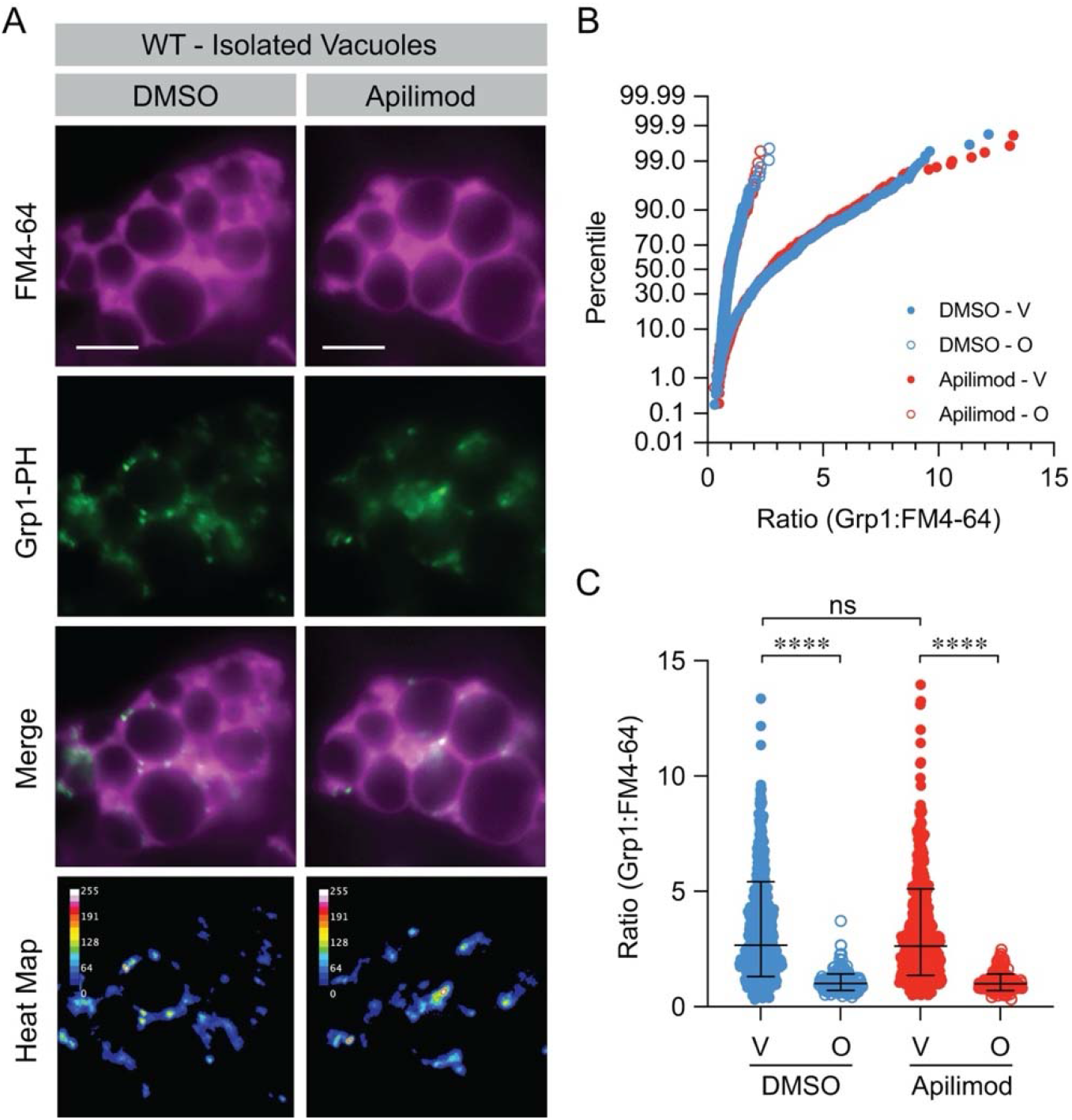
Apilimod has no effect on PI(3,4,5)P_3_ labeling. **(A)** Docking reactions using purified vacuoles were incubated for 30 min at 27°C with and labeled with GST-Grp1-PH as described. Reactions were treated with 150 µM Apilimod or DMSO and prepared for fluorescence microscopy. After incubation the reactions were place on ice and mixed with 5 µM FM4-64 to label entire vacuoles. Vacuoles were pelleted to remove excess antibody and resuspended in PS buffer. **(B)** Cumulative distribution plot of pooled experiments (n=3) (DMSO-V = 543; DMSO-O = 215; Apilimod-V = 503; Apilimod-O = 178). **(C)** Scatter plots of pooled data shown in **(B)**. Error bars represent geometric means ± geometric standard deviations. Significance was determined using one-way ANOVA for multiple comparisons and Tukey’s test for individual *p*-values. \*\*\*\**p*<0.0001; ns – not significant. Scale bars: 5 µm.

## Supplementary Information Detailed Computational Methods

### System construction

Vam7 PX domain was obtained from PDB 1KMD (Cheever et al., 2001). Residues 8–124 were included, and the N- and C- termini were capped with neutral ACE and CT3, respectively. WT and Y42A protein models were used to construct 4 conditions: WT–PI3P, WT–PIP3, Y42A– PI3P, and Y42A–PIP3. Membranes were generated with CHARMM-GUI and contained POPC:POPE:POPS:ergosterol:PIP (50:20:10:10:10). The target phosphoinositide was either PI3P (POPI13) or PI(3,4,5)P_3_ (POPI35). Two independent membrane builds were generated for each phosphoinositide composition. Each condition contained 4 simulation replicas. Replica 1 was used as the reference orientation for the PX domain. Replicas 2–4 employed branched orientations in which the protein was tilted by 45^◦^ around the *x* axis and rotated around the membrane-normal by 0°, 120°, or 240° This design sampled both lipid-packing and protein-orientation variabilities.

### HMMM model and membrane restraints

The atomistic membranes were converted to the highly mobile membrane mimetic (HMMM) representation using an in-house VMD/Tcl workflow. In the HMMM representation, lipid headgroups and upper acyl-chain segments are retained while the long hydrophobic tail regions are replaced by a short-chain organic solvent. This treatment accelerates lateral diffusion of lipids and enhances the sampling of peripheral membrane-binding events (Ohkubo et al., 2012; Hasdemir et al., 2026). Ergosterol was retained as full sterol.

During equilibration, harmonic restraints were applied to protein C_α_ atoms and select membrane atoms (*k* = 1.0 and 0.5 kcal mol⁻¹ Å⁻² during the first and second equilibration stages, respectively). During the production runs, only weak restraints (*k* = 0.05 kcal mol⁻¹ Å⁻²) were applied to select lipid atoms. A grid potential was also applied to the (SCSE) organic phase of the HMMM membrane to maintain its slab geometry while allowing the protein and lipid headgroups to sample the membrane.

### Simulation protocol

All systems were simulated in NAMD (Phillips et al., 2005, 2020). using CHARMM36m and CHARMM36-compatible lipid parameters. TIP3P water and KCl were used for solvation and ionization. Long-range electrostatics were calculated using particle-mesh Ewald (Darden et al., 1993). Nonbonded interactions used a 12 Å-cutoff, a 10-Å switching distance, and a 16 Å pair-list distance. Bonds involving hydrogen atoms were constrained.

Each system was minimized for 10,000 steps and equilibrated in 2 restrained steps. Production simulations were then performed for 250 ns per replica using a 2-fs timestep. Coordinates were saved every 50,000 timesteps (0.1 ns). The total production sampling was 4.0 *µ*s across the 16 replicas.

### Trajectory preparation

Production trajectories were wrapped using the membrane as the reference. Water and ions were removed from the wrapped trajectories to generate protein–membrane trajectories. The resulting trajectories were used for RMSD (root mean square deviation), depth, per-residue distance, UMAP (uniform manifold approximation and projection), and contact analyses.

### C**_α_** RMSD quality control

Protein C_α_ RMSD was calculated after least-squares fitting each trajectory frame to the first frame using C_α_ atoms. RMSD was used only as a quality-control metric and not for defining membrane binding. For a selection *S* containing *N_S_* C_α_ atoms, RMSD at time *t* was calculated according to

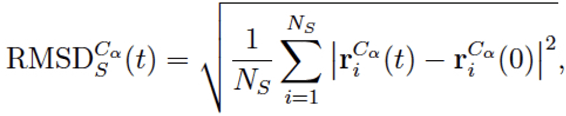

where **r***^C^ ^α^* (*t*) is the fitted position of atom *i* at time *t*.

### Membrane depth and stably bound intervals

For each frame, the closest protein atom along the membrane normal was compared with the plane formed by the phospholipid phosphorus atoms and the ergosterol O3 atom. The sign was assigned so that negative values indicate that the protein reaches or crosses the phosphate plane. A frame was considered as membrane-bound when

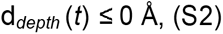

A stably membrane-bound interval was defined as a continuous bound segment lasting at least 30 ns. Since coordinates were saved every 0.1 ns, this definition included at least 300 consecutive membrane-bound frames.

### Per-residue membrane-distance profiles and UMAP analysis

For stably bound frame, the membrane-normal distance between each residue and the mem-brane phosphate plane was calculated. Residue-level distance profiles were averaged over stably bound frames for each replica and used to evaluate whether independent replicas sampled similar membrane-binding geometries. Stably bound frames were also represented by their per-residue membrane-distance profiles, standardized, and projected into two dimensions using UMAP (McInnes et al., 2018). DBSCAN was used only to organize dense regions of the stably bound ensemble and was not interpreted as defining distinct biological binding modes (Ester et al., 1996).

### Basic residue/PIP oxygen contact analysis

Contacts were calculated between closest Lys/Arg side-chain nitrogen atoms and phosphoinositide oxygen atoms using a 4.0 Å cutoff. The protein atoms included Lys NZ and Arg NE/NH1/NH2 atoms. The lipid atom set included all oxygen atoms of PI3P or PI(3,4,5)P_3_. The frame-wise number of unique basic residues contacting PIP was calculated by counting unique Lys/Arg residues with at least one side-chain nitrogen atom within 4.0 Å of a PIP oxygen atom.

---

**Table S1:** Membrane binding across simulation conditions. Stable intervals were defined as continuous membrane-bound segments lasting at least 30 ns. Values are reported as mean ± standard deviation across replicas where applicable.

| Condition | Replicas | Stably-bound fraction | Mean stable depth (Å) |
| --- | --- | --- | --- |
| WT-PI3P | 4/4 | $0.51 \pm 0.20$ | -3.79 |
| WT-PIP3 | 1/4 | $0.22 \pm 0.43$ | -4.14 |
| Y42A-PI3P | 1/4 | $0.20 \pm 0.41$ | -4.44 |
| Y42A-PIP3 | 1/4 | $0.19 \pm 0.37$ | -5.09 |

**Table S2:**
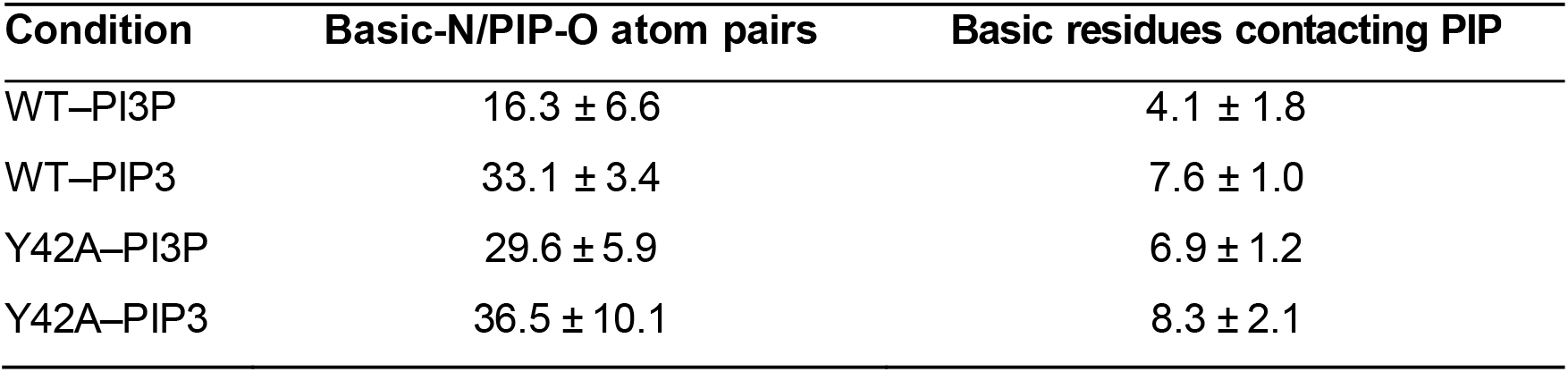
Basic residue/PIP contacts. Contacts were defined using a 4.0-Å cutoff between Lys/Arg side-chain nitrogen atoms and PI3P or PIP3 oxygen atoms. Values are mean ± standard deviation across replicas.

**Table S3.** Strains used in this study.

| **Strain** | **Genotype** | **Source** |
| --- | --- | --- |
| BJ3505 | *MATa pep4::HIS3 prb1-Δ1.6R his3-200 lys2-801 trp1Δ101 (gal3) ura3-52 gal2 can1* | (Jones et al., 1982) |
| DKY6281 | *MATα leu2-3,112 ura3-52 his3-Δ200 trp1-Δ901 lys2-801 suc2-Δ9 pho8Δ::TRP1* | (Klionsky and Emr, 1989) |
| CBP-Vam3 | *BJ3505, CBP-VAM3::Kan<sup>r</sup> nyv1Δ::nat<sup>r</sup>* | (Collins and Wickner, 2007) |
| SEY6210 | *MATα leu2-3,112 ura3-52 his3-Δ200 trp1-Δ901 lys2-801 suc2-Δ9* | (Klionsky and Emr, 1989) |
| BY4742 | *MATα, his3Δ1 leu2Δ0 lys2Δ0 ura3Δ0* | Open Biosystems (Huntsville, AL) |
| GFP-Ypt7 | *SEY6210, ypt7::HIS3 pRS304:GFP-Ypt7 (TRP1)* | (Wang et al., 2002) |
| Vps33-GFP | *DKY6281, VPS33-GFP* | (Wang et al., 2002) |
| *tep1Δ* | *BY4742, tep1Δ::G418<sup>r</sup>* | Horizon Discovery (Cambridge, UK) |
| *tsvps34* | *SEY6210, vps34Δ::TRP1, pRS416-VPS34TS* | (Stack and Emr, 1994) |
| AAY202 | *SEY6210, mss4Δ::HISMX6 YCplac111mss4<sup>ts</sup> (LEU2 CEN6 mss4<sup>ts</sup> 102)* | (Stefan et al., 2002) |
| mCh-Grp1-PH | *tsvps34, GST-mCHERRY-GRP1-PH (LEU1)* | This study |

